# Perturbing glycosylphosphatidylinositol (GPI)-anchor biosynthesis alters cell wall architecture and modulates fungal morphology

**DOI:** 10.64898/2026.06.05.730525

**Authors:** Hui Ting Chu, Isha Gautam, Surya Pavan Yenamandra, Tuo Wang, Prakash Arumugam

## Abstract

Filamentous fungi are widely used for industrial protein production but their tendency to form dense mycelial pellets limits nutrient transfer, oxygen uptake and fermentation efficiency. While cell wall components are known to influence fungal aggregation, the molecular mechanisms linking cell wall biosynthesis to macroscopic morphology remain poorly defined. Using *Aspergillus oryzae* as a model, we show that perturbation of glycosylphosphatidylinositol (GPI)-anchored protein biosynthesis influences fungal morphology by altering cell wall composition. Disruption of the GPI ethanolamine phosphate transferase 1 (mcd4) or inhibition of glucosaminyl phosphatidylinositol acyltransferase Gwt1 using antifungal drug manogepix (MGX) induced hyper-branching and weakened the cell wall. Notably, chemical inhibition of Gwt1 by MGX causes a marked transition from pelleted to dispersed mycelial growth. Solid-state NMR (ssNMR) analysis revealed reorganisation of galactosaminogalactan (GAG) and galactomannan (GM), including complete loss of cationic galactosamine (GalN), a key determinant of hyphal adhesion. Transcriptomic analysis revealed downregulation of genes involved in somatic cell fusion, linking altered wall composition to impaired germling aggregation. Strikingly, MGX treatment induced distinct morphological outcomes in other industrially relevant fungi, indicating species-specific cell- wall dependencies. Together, these findings identify GPI-anchored proteins as key regulators of fungal morphology and provide a framework for rational morphology engineering to improve industrial fermentation.

## Introduction

*Aspergillus oryzae* is an important filamentous fungus widely used in biotechnology for industrial production of recombinant proteins, enzymes and secondary metabolites ^1^ owing to its strong capabilities in protein expression, secretion and post-translational modification ^2^. However, the dynamic fungal morphology (compact pellets, loose mycelial clumps, or dispersed hyphae) in submerged culture remains a major challenge in controlling the outcome of large-scale fermentation, as it strongly influences culture viscosity and fermentation efficiency ^3,4,5^. Pelleted growth reduces viscosity but limits nutrients and oxygen diffusion into the pellet core, whereas dispersed mycelia enhance mass transfer but markedly increase viscosity, hindering gas-liquid exchange ^3^. The macro-morphology of filamentous fungi is influenced by inoculum concentration, pH, medium composition, agitation rate, etc ^6^. At a molecular level, pellet formation is thought to be largely governed by hydrophobic, electrostatic and specific interactions between spore wall components ^7^.

The *A. oryzae* cell wall is primarily composed of polysaccharides, such as α-glucan, β-glucan, chitin, galactomannan (GM) and galactosaminogalactan (GAG), and a small fraction of lipids and proteins (< 20%) ^8,9^. GAG and α-1,3-glucan mediate hyphal and conidial aggregation, respectively, in *A. oryzae* ^10^, while α-1,3-glucan promotes conidial aggregation in *A. fumigatus* ^11^. However, the precise mechanisms by which the cell wall composition affects mycelial aggregation remain to be fully elucidated.

Besides polysaccharides, glycosylphosphatidylinositol (GPI)-anchored proteins, galactomannoproteins and other surface proteins are also present in the fungal cell wall. GPI-anchors are glycolipids that post-translationally attach to the C-terminus of secretory proteins, tethering them to the plasma membrane or cell wall ^12^. GPI-anchored proteins play essential roles in ligand recognition, enzymatic activity, cell-cell interaction, and host defense^13^. Many participate in cell wall synthesis and remodeling, including the GEL family proteins (β-1,3-glucan elongation and branching) and the DFG family proteins (covalent attachment of GM to β-1,3-glucan-chitin core) ^13^. Moreover, GPI-anchors are involved in GM trafficking from Golgi body to cell wall, further highlighting their critical role in maintaining cell wall integrity ^14^. Thus, perturbation of GPI-anchor biosynthesis is expected to substantially affect cell-wall structure and composition.

GPI-anchor biosynthesis occurs in the ER and requires sequential action of more than 20 enzymes (**Supplementary Fig. 1**). One such enzyme is Mcd4 (morphogenesis checkpoint dependent) an integral membrane protein with 14 transmembrane segments ^15,16^, that was first identified in *Saccharomyces cerevisiae* mutant exhibiting defects in bud emergence and polarized growth ^15^. Mcd4 catalyses the transfer of phosphoethanolamine (EtN-P) to the first mannose (Man1) of phosphatidylinositol (PI) anchor, a conserved step in the GPI-anchor biosynthetic pathway^15,17,18^. *MCD4* deletion is lethal in *S. cerevisiae* ^19^, but the lethality can be rescued by expressing *Trypanosoma brucei* Gpi10p, which bypasses substrate requiring Mcd4 action ^20^. The *mcd4*-null studies revealed its role in ensuring proper tethering of GPI-anchored proteins to the β-1,6-glucan in the cell wall in *S. cerevisiae* ^20,21^. Furthermore, *mcd4* deletion alters cell wall composition, increasing chitin and chitin-linked alkali-insoluble β-1,6-glucan while reducing mannans ^22^. Importantly, Mcd4 supports normal ER function by mediating ATP uptake ^16^ and facilitating trafficking of GPI-anchored proteins from the ER to the Golgi ^20^.

Manogepix (MGX; APX001A) is an antifungal developed by Amplyx Pharmaceuticals that targets the GPI-anchor biosynthetic enzyme Gwt1 in fungi ^23^. Gwt1 works upstream of Mcd4 and catalyses the transfer of fatty acyl chains to inositol moiety of GPI anchor precursors. In *Candida albicans*, exposure to MGX at four times the minimal inhibitory concentration (MIC) resulted in depletion of cell-surface mannoproteins, increased cell size and elevated chitin content ^24^.

In this study, we perturbed GPI-anchor biosynthesis in *A. oryzae* by either deleting *mcd4* or by chemically inhibiting Gwt1 with MGX to investigate their effects on cell wall architecture, macro-morphology and recombinant protein secretion (**Figure 1a**). By integrating ssNMR and transcriptomics analyses of the perturbed strains, we identified potential morphology-related genes that may serve as targets for rational morphology engineering.

**Figure 1:**
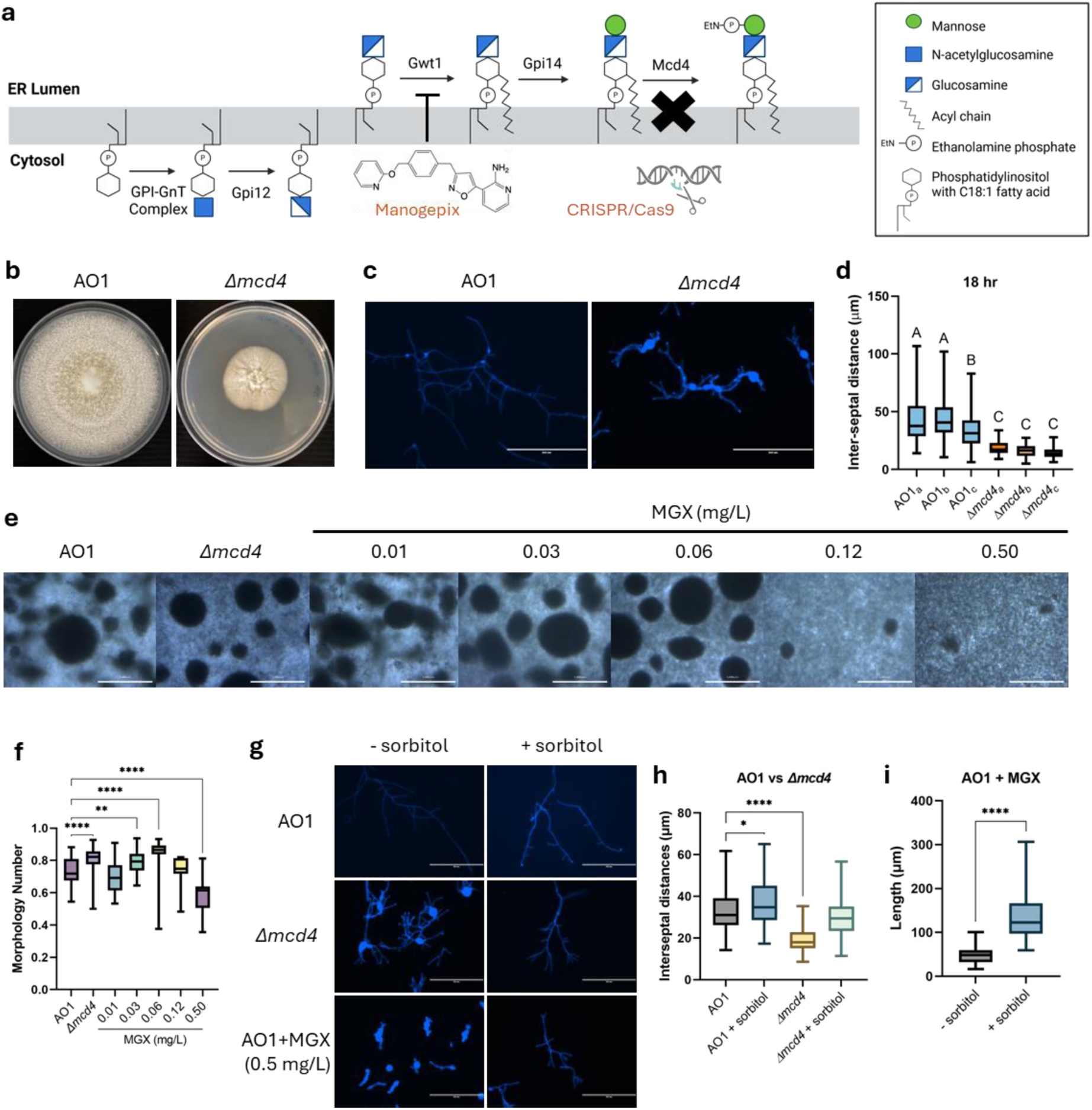
Perturbation of GPI-anchor biosynthesis alters fungal morphology and affects hyphal integrity in *A. oryzae*. (a) Schematic of GPI-anchor biosynthesis at the endoplasmic reticulum (ER) lumen and strategies to disrupt the pathway through *mcd4* deletion and Gwt1 inhibition via manogepix (MGX). Created with Biorender.com and ChemDraw v21.0.0. (b) Colony morphology of AO1 and *Δmcd4* strains grown on potato dextrose agar (PDA) for 7 d at 30 °C. (c) Hyphal morphology of AO1 and *Δmcd4* strains in *Aspergillus* minimal medium (AMM) supplemented with 1% glucose for 18 h. Septa were stained with calcofluor white (CFW) to visualize inter-septal distance. Scale bar, 200 µm. (d) Inter-septal distances of hyphae from ImageJ analyses of CFW-stained AO1 and *Δmcd4* hyphae. Statistical significance was determined by one-way ANOVA with Tukey’s multiple comparisons test using GraphPad Prism v9.3.1; datasets labelled with different letters differ significantly at P < 0.05. (e) Macro-morphology comparison of AO1, *Δmcd4* and MGX-treated AO1 at increasing concentrations of MGX grown in 5×DPY medium for 4 d. Scale bar, 1000 µm. (f) Morphology number of fungal pellets (n > 10) from ImageJ analyses of AO1, *Δmcd4* and MGX-treated AO1 microscopy images. (g) Sorbitol partially rescues the phenotype of *Δmcd4* and AO1 + MGX strains grown in AMM without (–) or with (+) 1.2 M sorbitol. Cells were stained with CFW. Scale bar, 200 µm. (h) Inter-septal distances and (i) cell length of hyphae from ImageJ analyses of CFW-stained hyphae microscopy images (n > 10). Statistical significance was determined by one-way ANOVA with Dunnett’s multiple comparisons test against AO1 or Student’s t-test using GraphPad Prism v9.3.1: P ≤ 0.05 (*), P ≤ 0.01 (**), P ≤ 0.001 (***), P ≤ 0.0001 (****).

## Results

### Perturbation of GPI anchor synthesis affects cell wall integrity and morphology

The GPI-anchored proteins in *A. oryzae* RIB40 were predicted *in silico* by screening the complete proteome on UniProt and FungiDB using SignalP-6.0 and NetGPI-1.1, identifying 78 candidates (**Supplementary Table 1**). Many of these GPI-APs are implicated in cell wall polysaccharide remodelling or mediate interactions between fungal cells ^25^. We therefore investigated whether perturbation of GPI anchor biosynthesis influences pellet morphology. We examined the phenotypic consequences of deleting the *mcd4* and *gwt1* genes which encode the GPI ethanolamine phosphate transferase 1 and glucosaminyl phosphatidylinositol acyltransferase, respectively ^15,26^. *A. oryzae* strain RIB40 Δ*wA*::amyB-Lys-Arg-HLY (hereafter termed AO1), which expresses human lysozyme (HLY) as a secretion reporter ^27^, was used as the base strain for our experiments. Both *Δmcd4* and *Δgwt1* mutants displayed markedly altered morphology with fewer hyphae and slower growth (**Figure 1b**). However, the *Δgwt1* mutants could not be successfully revived from glycerol stocks, preventing further characterization (data not shown). The *Δmcd4* mutant exhibited smaller colony radius and diminished conidiation compared to AO1 on potato dextrose agar (PDA) (**Figure 1b**). It also displayed a hyper-branching phenotype as reflected by significantly shorter inter-septal distance relative to AO1 (**Figure 1c, d**). Hyper-branching is common in fungal strains with weakened cell walls, as reduced inter-septal distance increases cell rigidity ^28^. Consistent with this possibility, the *Δmcd4* mutant showed hyphal swelling and localized cell wall rupture, indicating compromised cell wall integrity (**Supplementary Fig. 2a)**.

Owing to the poor revivability of the frozen Δ*gwt1* stocks, we explored a chemical genetic strategy using MGX, a Gwt1 inhibitor, which enables dose-dependent and reversible inhibition of protein function ^29,23^. We treated the AO1 fungal cultures with varying doses of MGX (0.01–0.50 mg/L) and monitored their morphology over the course of 4 days. While low MGX concentration-treated fungal cultures largely formed pellets, MGX treatment at 0.12 and 0.50 mg/L resulted in largely dispersed mycelia with a few small pellets (**Figure 1e**). This was accompanied by a significant decrease in morphology number in the 0.50 mg/L MGX-treated AO1 culture (**Figure 1f**). The *Δmcd4* mutant, however, formed spherical pellets similar to the AO1 strain (**Figure 1e**).

Swollen hyphae, indicative of cell wall stress, were also observed in AO1 strain treated with 0.50 mg/L MGX (**Supplementary Fig. 2b**). Consistent with this observation, the addition of 1.2 M sorbitol to the *Δmcd4* mutant and 0.50 mg/L MGX-treated AO1 partially rescued the cell wall defect by reducing hyphal swelling and restoring normal branching behaviour in *Δmcd4* mutant (**Figure 1g,h**) and significantly increased the cell length of MGX-treated AO1 (**Figure 1i**). However, sorbitol addition did not rescue the pellet morphology in 0.50 mg/L MGX-treated AO1 (**Supplementary Fig. 2c**), suggesting that defective GPI-anchor synthesis continues to influence macro-morphology even after sorbitol addition.

### MGX modulates A. oryzae morphology through Gwt1 inhibition and is the most effective at the conidial stage

The chemical perturbation of Gwt1 and deletion of *mcd4* produced different morphological outcomes, despite both strategies targeting the GPI anchor synthesis. To test whether the morphological effects induced by MGX were due to Gwt1 inhibition, we introduced the MGX-resistance mutations previously identified in *S. cerevisiae* Gwt1 into the *A. oryzae* AO1 *gwt1* gene ^30^. Mutations in ScGwt1 G132, F238, S170, V168, L136, I141, F171, Y400 and Y408 resulted in resistance towards MGX ^30^. Sequence alignment of ScGwt1 and AoGwt1 shows that the residues I141, V168, F171, and F238 in ScGWT1 are conserved and correspond to I150, V177, F180, and F248 in AoGwt1, respectively (**Figure 2a**) Structural alignment of AoGwt1 ^31^ with the ScGwt1–MGX complex (PDB ID: 8XIK) ^30^ yielded a backbone root-mean-square-deviation (RMSD) of 1.378 Å across 341 aligned residues, confirming conservation of these sites within the MGX binding pocket (**Figure 2b**). We introduced the MGX-resistance mutations in AoGwt1 via CRISPR-mediated gene editing ^32^, and compared the sensitivity of AO1, Gwt1-I150A and Gwt1-V177A+F180A to MGX. Gwt1-V177A+F180A showed enhanced resistance towards MGX, while Gwt1-I150A showed partial resistance up to 0.25 mg/L (**Figure 2c**). We then compared the morphology-modulating effects of MGX on wild type and MGX-resistant strains in submerged cultures. As observed previously, MGX treatment (0.50 mg/L) of AO1 resulted in predominantly dispersed mycelia with significant reduction in morphology number (**Figure 2d,e**). In the partially resistant Gwt1-I150A mutant, we observed a mixed morphology consisting of both dispersed hyphae and pellets, resulting in a huge range of morphology number (**Figure 2d,e**). Importantly, the MGX-resistant Gwt1-V177A+F180A mutant strain retained its pellet morphology and was indistinguishable from its DMSO-treated control, with no significant difference in morphology number (**Figure 2d,e**). These results demonstrate that MGX exerts its morphological effects through by inhibition of Gwt1 in *A. oryzae*, rather than through off-target toxicity.

**Figure 2:**
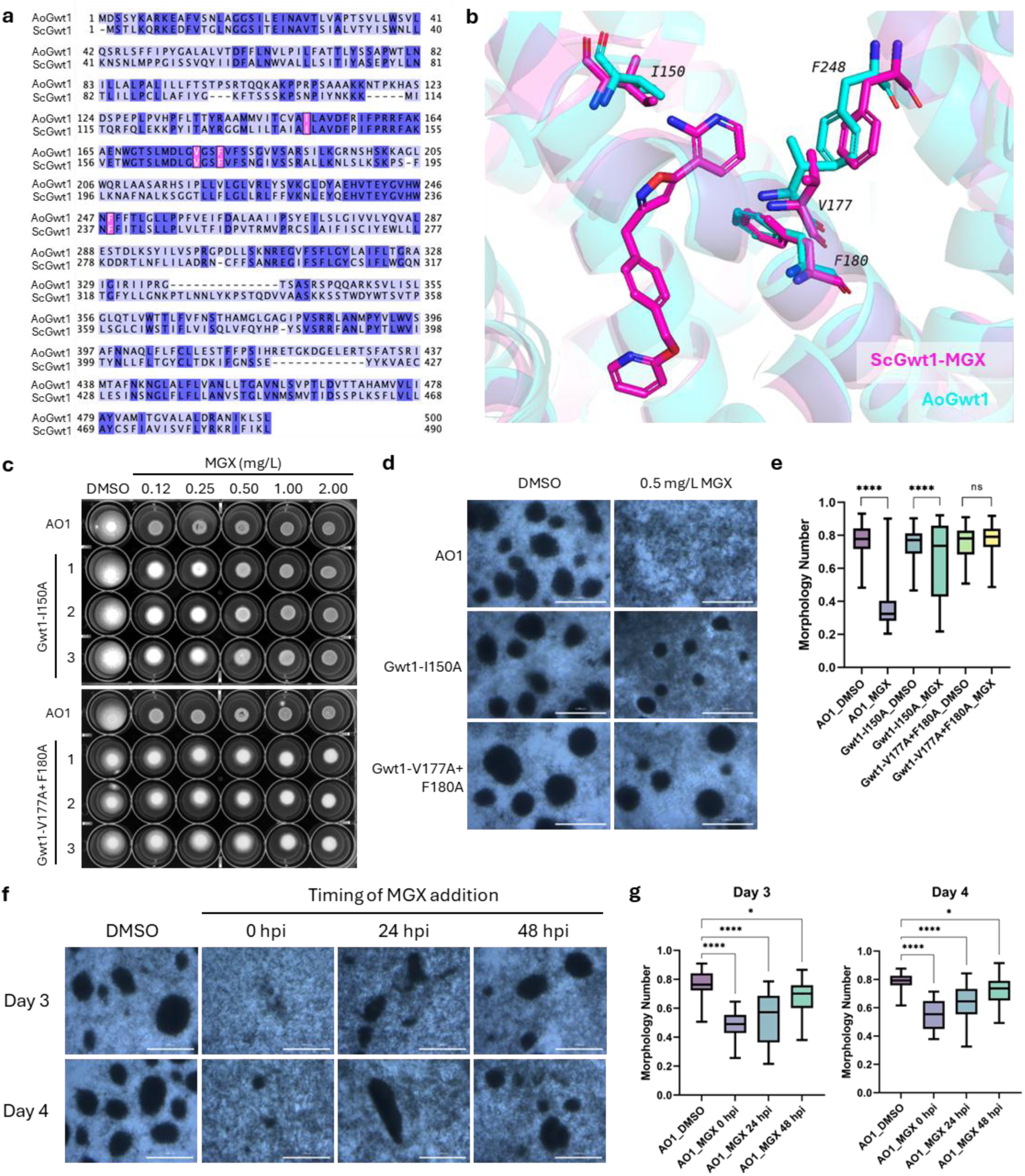
MGX perturbs *A. oryzae* morphology through Gwt1 inhibition. (a) Sequence alignment of *S. cerevisiae* Gwt1 (ScGwt1) and *A. oryzae* Gwt1 (AoGwt1). Residues I150, V177, F180, and F248 in *A. oryzae* (pink) are homologous to MGX-resistance sites I141, V168, F171, and F238 in *S. cerevisiae*. (b) Structural superimposition of AoGwt1 (cyan) and ScGwt1−MGX complex (magenta, PDB: 8XIK). (c) MGX minimal inhibitory concentration (MIC) on AO1 and alanine-substituted mutants (Gwt1-I150A and Gwt1-V177A+ F180A) grown on PDA for 2 days (d) Macro-morphology comparison of AO1 and alanine-substituted mutants following 0.50 mg/L MGX treatment in 5×DPY medium compared to DMSO control for 3 d. Scale bar, 1000 µm. (f) Time-resolved effects of MGX (0.50 mg/L) on AO1 morphology against DMSO control. MGX was dosed at 0, 24 and 48 h post-inoculation (hpi) and macro-morphology of fungal culture analysed after 3-4 d. Scale bar, 1000 µm. Morphology number of fungal pellets (n > 10) from ImageJ analyses of fungal pellet microscopy images for (e) alanine-substituted mutants and (g) time-resolved MGX treatment against AO1 (DMSO control). Statistical significance was determined by one-way ANOVA with Dunnett’s multiple comparisons test against AO1 using GraphPad Prism v9.3.1: P ≤ 0.05 (*), P ≤ 0.01(**), P ≤ 0.001 (***), P ≤ 0.0001 (****).

Pellet formation in fungal cultures can result from aggregation of either conidia or mycelia or via a combination of both ^4,33^. We therefore compared the morphology-modulating effects of MGX on conidia and mycelia by adding 0.50 mg/L MGX at 0, 24 and 48 h post-inoculation (hpi) and analysed their effects on morphology on days 3 and 4 (**Figure 2f**). The AO1 culture treated with MGX 0 hpi resulted in the most pronounced morphological changes, yielding predominantly dispersed mycelia morphology and significantly reduced morphology number (**Figure 2g**). While the AO1 culture treated with MGX at 24 and 48 hpi resulted in some pellet formation, these cultures still exhibited higher proportion of dispersed mycelia compared to DMSO-treated control. The pellet analyses showed significant reduction in morphology number, although the difference from DMSO control is less significant in 48 hpi MGX-treated culture compared to 0 and 24 hpi MGX-treated culture (**Figure 2g**). These observations indicate that MGX can disrupt the GPI-anchor biosynthesis to affect morphology in both spores and actively growing mycelia but is more effective when added at conidial stage.

### Morphological modulation by chemical inhibition of Gwt1 and mcd4 deletion uncovers opposing effects on growth and secretion

To assess whether perturbation of GPI-anchored proteins and morphology influences protein secretion, we compared HLY production in AO1, Δ*mcd4* and AO1 treated with 0.5 mg/L MGX (AO1 + MGX), over 3–5 days of submerged fermentation. As expected, the *Δmcd4* mutant formed pellets similar to AO1 and the MGX-treated AO1 exhibited dispersed mycelial morphology (**Figure 3a,b**). Despite exhibiting similar morphology and biomass-normalised culture viscosity as AO1 (**Figure 3a,c**) the biomass accumulation in *Δmcd4* was consistently lower than that of AO1 (**Figure 3d**), reflecting a growth defect. The biomass-normalized HLY titres was, however, comparable to AO1 on days 3 and 4, and exceeded AO1 by day 5, indicating that its secretion efficiency was not affected by the growth defect and the secreted protein/ biomass ratio remains unchanged (**Figure 3e**).

**Figure 3:**
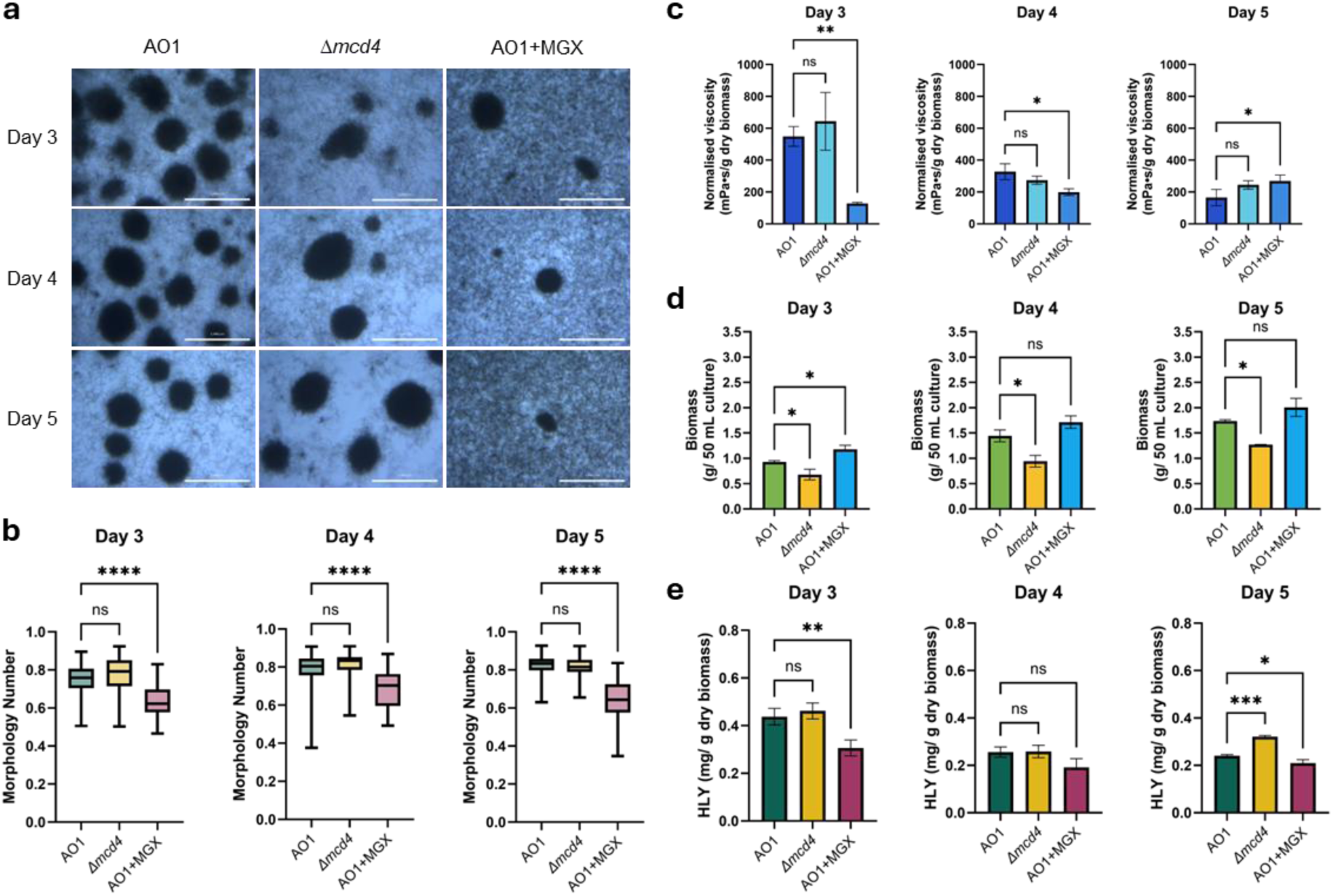
Genetic and chemical perturbation of GPI-anchor biosynthesis differentially affect morphology, growth, viscosity, and HLY secretion. (a) Macro-morphology comparison of *A. oryzae* AO1, *Δmcd4* and AO1 + MGX (0.5 mg/L MGX) cultured in 5×DPY for 3-5 d. Scale bar, 1000 µm. (b) Morphology number of fungal pellets from ImageJ analyses of *A. oryzae* pellet microscopy images. (c) Biomass-normalised viscosity, (d) dry biomass, and (e) biomass-normalised HLY of the *A. oryzae* cultures. Data represent mean ± s.d. of three biological replicates cultured in independent flasks. Statistical significance was determined by one-way ANOVA with Dunnett’s multiple comparisons test against WT using GraphPad Prism v9.3.1: P ≤ 0.05 (*), P ≤ 0.01(**), P ≤ 0.001 (***), P ≤ 0.0001 (****).

The freely dispersed morphology of mycelia in MGX-treated strain was associated with both reduced viscosity (**Figure 3c**) and enhanced biomass accumulation (**Figure 3d**). However, these apparent advantages did not translate into improved protein secretion. Biomass-normalized HLY titres decreased by 30% and 13 % on days 3 and 5, respectively (**Figure 3e**), indicating that the secretion efficiency was partially impaired by GPI-anchor biosynthesis inhibition. Although the volumetric enzymatic production in MGX-treated AO1 culture showed significant reduction on day 3, it remained largely comparable to that of AO1 on days 4 and 5 (**Supplementary Fig. 3**). These results suggest that MGX treatment effectively lowers *A. oryzae* culture viscosity and promotes biomass accumulation, while maintaining overall volumetric protein secretion at levels comparable to AO1.

### Solid-state NMR and transcriptomics data revealed reduced cell wall GAG and GM levels in response to perturbation of GPI-anchor biosynthesis

Since GPI-anchored proteins play important roles in cell wall biosynthesis, assembly and maintenance, we examined how perturbation of GPI-anchor biosynthesis alters cell wall polysaccharide organisation. The cell wall composition of AO1, *Δmcd4* and AO1 + MGX were analysed using ssNMR spectroscopy. This technique enables direct characterisation of intact, insoluble cell walls, providing atomic-level insights into the native polysaccharide composition, organization, and dynamics ^34,35^.

Dynamically distinct structural domains were selectively probed using different polarization techniques: the hydrophobic and rigid core was probed using ^1^H-^13^C cross-polarisation (CP)- based 2D CORD experiments ^36,37^, while the hydrated and mobile outer shell was examined using through-bond ^13^C connectivity tracked by direct polarisation (DP)-based 2D J-INADEQUATE experiments ^38,39,40^. The rigid core of the *Δmcd4* mutant cell wall was largely comparable to that of AO1, with only a slight increase in type-a α-1,3-glucan (A^a^), while type-b (A^b^) remained unchanged. Interestingly, redistribution of chitin allomorphs was also observed, where a decrease in type-a chitin (Ch^a^) was compensated by an increase in Ch^b^, while β-1,3-glucans (B) levels were relatively stable (**Figure 4a,b** and **Supplementary Table 2**). In contrast, in the MGX-treated AO1 strain, a reduction of linear β-glucan chain in the inner core was accompanied by an increase in Ch^b^ and A^b^ (**Figure 4a,b** and **Supplementary Table 2**). These changes reflect a compensatory response, in which key structural organizational components are offset by increases in rigid polysaccharides such as α-glucan and chitin ^41^. The increase in chitin content helps to enhance the cell wall stability and rigidity and represents a conserved adaptive mechanism observed across many fungal species exposed to cell wall stress ^42,43^.

**Figure 4:**
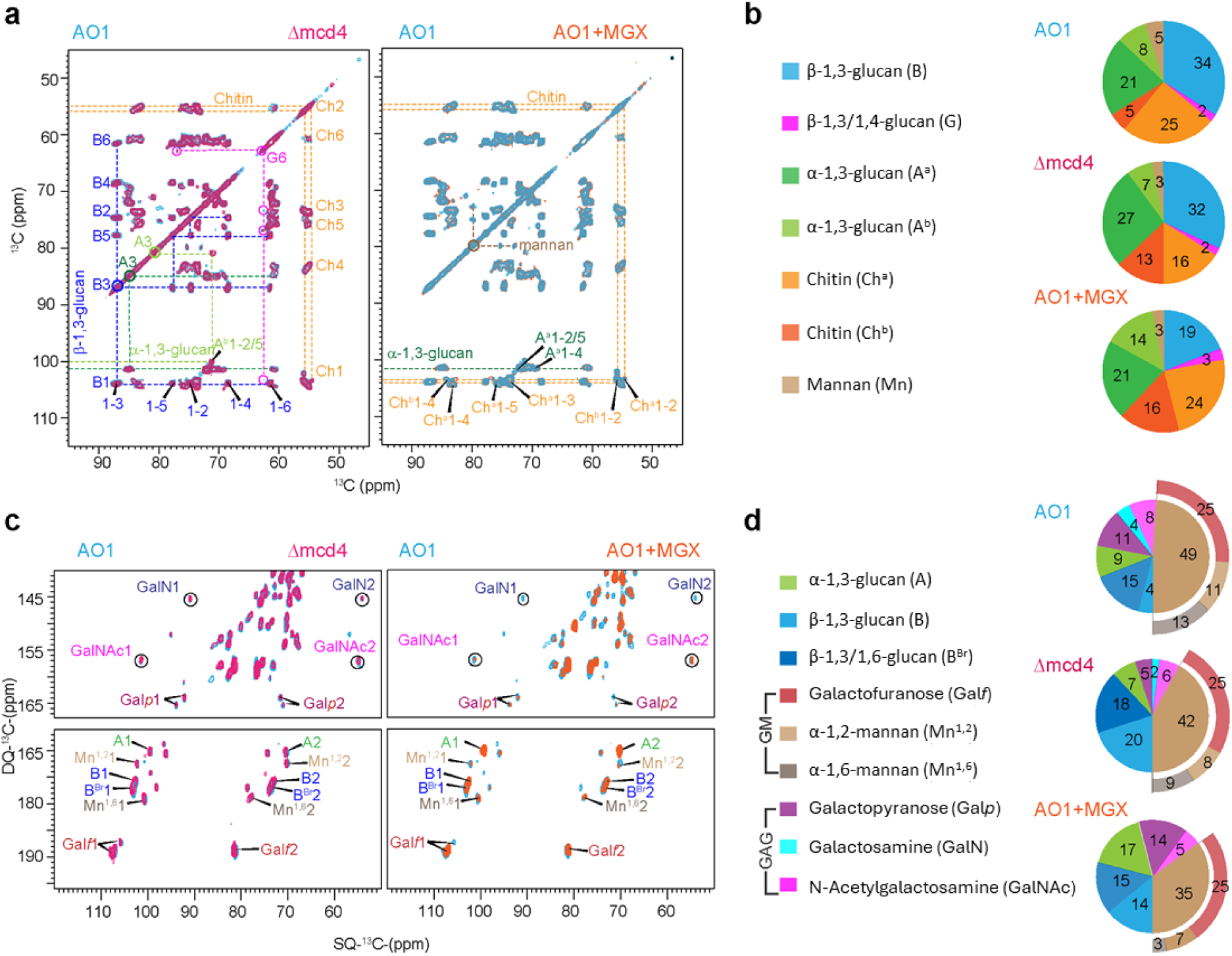
Perturbation of GPI-anchor biosynthesis alters the cell wall organization of *A. oryzae*. a) Overlay of 2D ^13^C-^13^C CORD spectra comparing AO1, *Δmcd4* and MGX-treated AO1 in cyan, magenta and orange, respectively. The spectra highlight changes in rigid polysaccharides, including the emergence of a distinct α-1,3-glucan form upon drug treatment, accompanied by increased chitin content. Colour coding and abbreviations are consistent throughout α-1,3-glucan (A), β-1,3-glucan (B), and chitin (Ch). Cross-peaks are indicated by dashed lines, including correlations originating from the diagonal resonances. (b) Relative abundance of rigid polysaccharides with consistent colour coding. (c) Overlay of 13C-13C DP INADEQUATE spectra highlighting the mobile cell wall components, using the same colour schemes. Mobile polysaccharides include α-1,3-glucan (A), β-1,3-glucan (B), β-1,3/1,6-glucan (B^Br^), galactofuranose (Gal*f*), α-1,2-mannan (Mn^1,2^), α-1,6-mannan (Mn^1,6^), galactopyranose (Gal*p*), galactosamine (GalN), N-acetylgalactosamine (GalNAc), with consistent colour coding and abbreviations and colour coding. These data reveal compositional changes in the outer cell wall: a decrease in GM components and absence of GalN in MGX-treated AO1. (d) Molar composition of mobile cell wall polysaccharides estimated from resolved spin pairs in 2D ^13^C DP INADEQUATE spectra showing reduced GM and GAG under GPI-anchor-perturbed conditions.

In the hydrated and mobile outer shell, both *Δmcd4* and MGX-treated AO1 showed a decline in the content of GAG and GM, accompanied by a compensatory increase in α-glucan and β-glucan (B or B^Br^) (**Figure 4c,d** and **Supplementary Table 2**). This selective depletion can be directly linked to the role of GPI anchors in polysaccharide trafficking and incorporation. GM is synthesized in the Golgi and transported to the cell surface via GPI-linked intermediates that are directly attached to the mannan chain ^14^. This explains why disruption of GPI anchor biosynthesis preferentially reduces mannan components (α-1,2- and α-1,6-linked mannans), while galactofuranose (Gal*f*) side chains are largely retained. The interaction between GM and β-1,3-glucan in the cell wall plays an important role in proper cell wall synthesis by providing cell wall support. Inhibition of GM synthesis disrupts cell wall organization, leading to a characteristic phenotype of hyperbranched mycelium and swollen filaments ^44–47^, as also observed in GM-depleted AO1 (**Supplementary Fig. 2d,e**).

More importantly, the cationic galactosamine (GalN) units of GAG were completely lost in MGX-treated AO1, while N-acetylgalactosamine (GalNAc) and galactopyranose (Gal*p*) were retained (**Figure 4d**). A similarly restructured GAG composition has recently been observed in *Aspergillus sydowii* under hypersaline conditions, and this change likely disrupts GAG-mediated surface adhesion and mycelial aggregation ^48^. Consistent with this, the absence of GalN correlates with the observed dispersed mycelial morphology, highlighting its functional role in the hyphal aggregation and biofilm formation. While the levels of branched beta-glucan (B^Br^) were relatively unchanged, both Δ*mcd4* mutant and MGX treatment induced redistribution towards more linear β-glucan chains. This suggests that the GPI-anchor perturbation affects not only composition but also the structure and organization of polysaccharides.

### Transcriptomics analyses revealed differential response to GPI-anchor perturbation induced through genetic and chemical genetic approaches

The cellular responses to GPI-anchor biosynthesis perturbation were investigated by the transcriptomic profiling of AO1, Δ*mcd4* and MGX-treated AO1 strains cultured in 5×DPY medium for 3–4 days (**Figure 5a**). Among the 12,074 genes in the *A. oryzae* RIB40 genome, 913 and 893 differentially expressed genes (DEGs; |log2 fold change (log_2_FC) | > 1.5, FDR < 0.05) were identified in *Δmcd4* mutants on days 3 and 4, respectively, whereas 2,183 and 2,348 DEGs were detected in MGX-treated AO1 cells (**Figure 5b**). MGX treatment induced more pronounced transcriptomics changes, as reflected by with the broader range of log_2_FC values (**Supplementary Fig. 4**). Heatmap analysis revealed distinct DEG clusters with shared or condition-specific expression patterns (**Supplementary Fig. 5**). Principal component analysis (PCA) showed the biological triplicates clustered together, with clear separation among the four treatment groups on day 3. However, AO1 and *Δmcd4* showed reduced separation on day 4 (**Supplementary Fig. 6**). This less distinct transcriptomic profile between AO1 and *Δmcd4* may explain the similarities in their morphologies. RT-qPCR analysis of selected upregulated and downregulated genes further validated the RNA-seq data, showing consistent expression trends (**Supplementary Fig. 7**).

**Figure 5:**
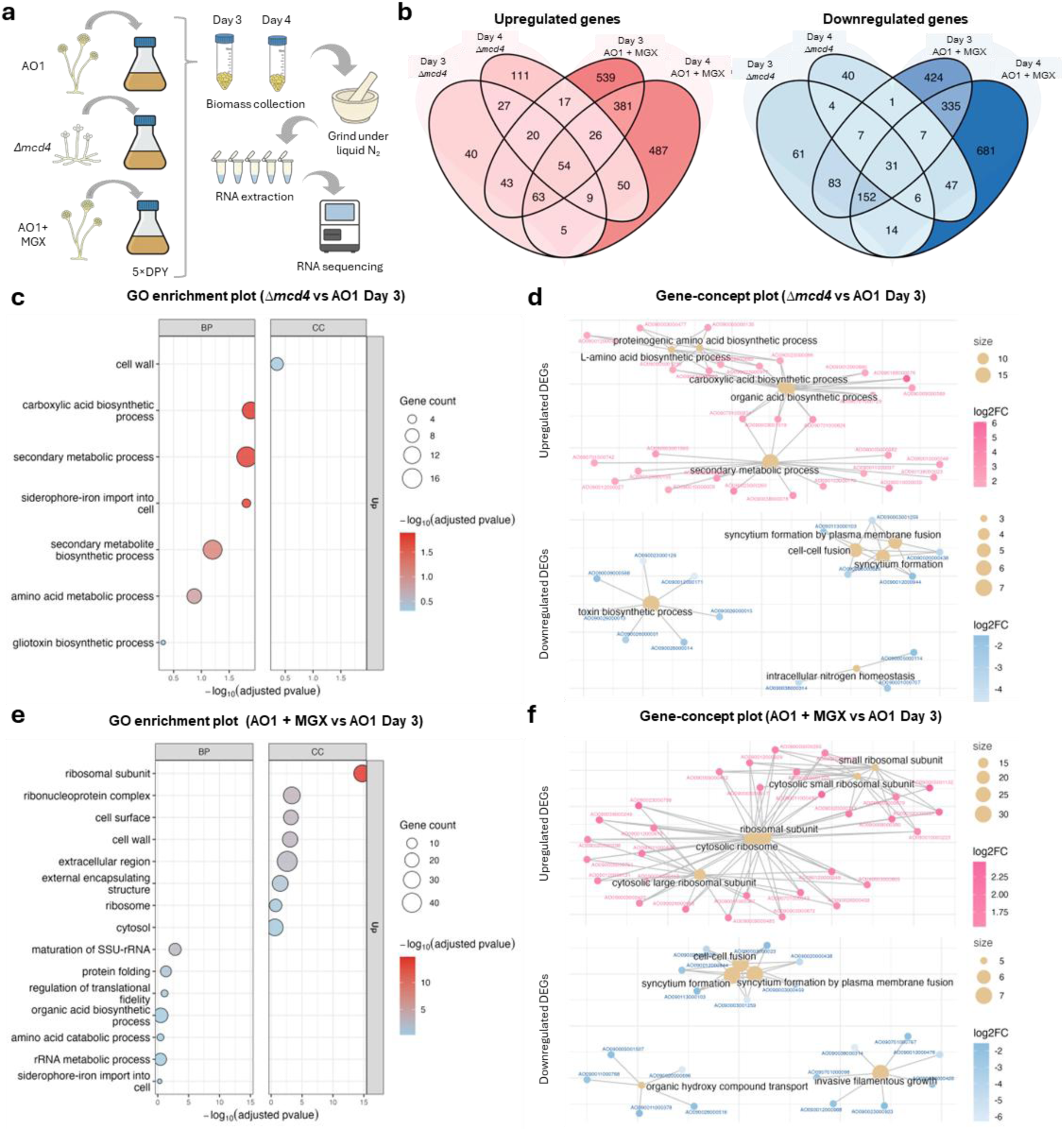
Transcriptomics profiling reveals widespread differential gene expression during perturbation of GPI-anchor biosynthesis. (a) Schematic diagram of workflow for biomass preparation and RNA extraction (Icons generated by ChatGPT 5, OpenAI, 16 September 2025). (b) Venn diagram of DEGs identified in *Δmcd4* and MGX-treated AO1 on days 3 and 4 relative to AO1, showing that MGX-treatment induced stronger transcriptomics response. GO enrichment plot from over-representation analysis (ORA) conducted on DEGs identified in (c) *Δmcd4* and (e) MGX-treated AO1 relative to AO1 on day 3. Enriched GO terms were filtered by REVIGO and GO terms with dispensability < 0.5 are displayed in bubble plot. Bubble size represents the gene number while the colour reflects the p-value. Gene–concept networks highlighting DEGs associated with multiple enriched GO terms in (d) *Δmcd4* and (f) MGX-treated AO1 relative to AO1 on day 3. Lines connect shared genes among terms. Each beige node represents a GO term, and node size reflects the number of associated genes. The colour of the gene nodes represents the log_2_FC.

Over-representation analysis (ORA) was conducted to identify GO terms enriched by the up-and down-regulated genes in *Δmcd4* and MGX-treated AO1. GO terms shortlisted by REVIGO with dispensability < 0.5 were plotted in the GO enrichment bubble plots (**Figure 5c,e** and **Supplementary Fig. 8a,b**). GO terms enriched by downregulated genes were excluded by REVIGO in most cases, except in MGX-treated AO1 on day 4 (**Supplementary Figure 8b**). Gene-concept network plots highlighted specific cluster of genes driving the enriched GO terms (**Figure 5d,f** and **Supplementary Fig. 9a,b**).

The analyses revealed distinct pathway enrichment profiles in genetic and chemical-genetic perturbation of GPI-anchor biosynthesis. The Δ*mcd4* mutant upregulated the secondary metabolite, amino acid and carboxylic acid biosynthetic pathways, while MGX treatment upregulated genes associated with protein synthesis such as ribosomal subunit, protein folding, regulation of translational fidelity, and small subunit ribosomal RNA (SSU-rRNA). GPI-anchor perturbation induced transcriptional changes associated with the cell wall remodelling in both conditions. However, GO enrichment analysis revealed stronger enrichment in MGX-treated AO1 (−log_10_(adjusted p-value) = 3.06 on day 3) compared to *Δmcd4* (**Figure 5c,e** and **Supplementary Fig. 8b**), suggesting more extensive cell wall remodelling upon chemical perturbation. By contrast, genes associated with siderophore-iron import into cell were enriched among upregulated genes in both conditions, with a stronger enrichment in *Δmcd4* (−log_10_(adjusted p-value) = 1.81 on day 3). This upregulation of iron homeostasis related pathways may reflect oxidative stress under GPI-anchor perturbation, particularly in *Δmcd4*, as heme is an essential cofactor for detoxification enzymes, including catalase and peroxidases^49^.

Upregulation of ribosomal proteins is typically associated with increased protein synthesis, either during rapid growth or as part of a cellular stress response to restore translation ^50^. Given that GPI-anchor biosynthesis occurs in the ER, disrupting this pathway is expected to accumulate misfolded GPI-anchored proteins intermediates, thereby inducing ER stress ^51,52^. Consistent with this, MGX-treated AO1 exhibited upregulation of multiple components of unfolded protein response (UPR) and ER-associated degradation (ERAD). The molecular chaperone BipA (AO090003000257) was strongly upregulated (log_2_FC = 3.2), alongside several unfolded protein-binding proteins (AO090012000995, AO090011000764, AO090102000620, AO090011000513, AO090103000007; log_2_FC = 1.5 to 2.4), indicative of an activated proteostasis network. On the other hand, amino acid biosynthesis was upregulated in *Δmcd4*, a response associated with late stage ER stress response ^53^. Despite a pronounced transcriptional signature of ER stress, both the *Δmcd4* and AO1 + MGX strains displayed tunicamycin tolerance comparable to AO1, with all strains tolerating up to 10 µg/mL (**Supplementary Fig. 10**). Although protein secretion in MGX-treated AO1 was reduced by 30% on day 3, it recovered to levels comparable to AO1 (**Figure 3e**). Notably, MGX-treated AO1 showed increased biomass accumulation (**Figure 3d**). Together, these results suggest that despite ER stress induction following GPI-anchor disruption, cellular proteostasis remains sufficiently buffered to sustain growth and secretion. GPI-anchor-perturbed strains exhibited cell wall stress, as evidenced by swollen cells and hyper-branched mycelia, which were partially rescued by sorbitol (**Figure 1g**). This stress likely activates the UPR and cell wall integrity (CWI) pathway to promote cell wall repair and restore cellular homeostasis ^54–56^. Notably, *Δmcd4* and MGX-treated AO1 appeared to engage distinct adaptive responses to overcome the cell wall stress. In *Δmcd4* mutant, the upregulation of amino acid, secondary metabolite, polyketide and siderophore biosynthesis may help to mitigate intracellular oxidative stress ^57,58^. By contrast, AO1 + MGX responded to cell wall stress through active cell wall remodelling by engaging genes associated with fungal-type cell wall and genes encoding ribosomal proteins. Together, these findings suggest that *Δmcd4* and MGX-treated AO1 experience and respond to GPI-anchor perturbation differently. The mutant may have undergone longer-term adaptation to chronic defect in GPI- anchor biosynthesis, whereas acute inhibition by MGX imposes a more immediate stress that requires active cell wall remodelling for survival.

Although many of the GO terms enriched by downregulated genes were excluded in REVIGO, the gene-concept map showed GO terms related to syncytium formation by plasma membrane fusion, syncytium formation and cell-cell fusion being enriched among the same cluster of downregulated genes in *Δmcd4* and MGX-treated AO1 on day 3 (**Figure 5d,f**). In filamentous fungi, somatic cell-cell fusion underpins genetic exchange, nutrient sharing and colony aggregation ^59,60^. Although less frequent in asexual species, conidial anamostosis tubes (CAT)-mediated fusion has been reported in *A. oryzae*, promoting germling fusion events ^61^. Consistently, homologues of *Neurospora crassa* fusion genes in *A. oryzae* were significantly downregulated under GPI-anchor perturbation conditions (**Supplementary Table 3**). Moreover, MGX addition prior to germination produced the most pronounced dispersed mycelia morphology (**Figure 2e)**, implicating early fusion events promoting hyphal aggregation in untreated culture. Notably, the ortholog of *N. crassa ham-7* (AO090020000438) encoding a GPI-anchored protein, was strongly repressed in MGX-treated AO1 (log_2_FC = −4.5 to −5), suggesting a potential link between GPI-anchor perturbation and impaired fusion. Together, these findings nominate cell-cell fusion as a previously underappreciated determinant of fungal macro-morphology and pellet formation.

### Correlation between ssNMR-defined structural changes in cell wall and transcriptomic changes elicited by perturbation of GPI-anchor biosynthesis

To assess whether transcriptional regulation of polysaccharide biosynthetic genes could explain the ssNMR-defined changes in cell wall structure, we analysed the expression of genes involved in α-glucan, β-glucan, chitin, GAG, GM and GalN biosynthesis (**Supplementary Fig. 11a**). α-Glucan biosynthesis genes (*agsB*, *agsC*, *amyD* and *amyG*) were broadly upregulated, consistent with increased α-glucan abundance in the mobile phase of *Δmcd4* and in both the core and mobile phases of MGX-treated AO1. Likewise, downregulation of GAG biosynthesis genes (*gtb3*, *agd3* and *ega3*) correlated with reduced GAG levels in both strains. Notably, *agd3* encoding GalNAc deacetylase was downregulated in MGX-treated AO1 (log_2_FC = −1.5 on day 4), potentially accounting for the complete absence of GalN in the MGX-treated AO1. Reduced expression of *ktr4* encoding α-1,2-mannosyltransferase was also consistent with the decreased level of mannans.

While we observed a good correlation between compositional changes in α-glucan, GAG, GM and GalN and the expression of the corresponding biosynthetic genes, there was no such link observed for β-glucan and chitin levels. Regulation of genes involved in β-glucan biosynthesis was heterogeneous, with mixed transcriptional changes and no significant alteration in the expression of *fksA* encoding β-1,3-glucan synthase. Despite minimal changes in the expression of chitin synthase genes, the chitin content increased following perturbation of GPI-anchor biosynthesis. This suggests that post-transcriptional or other regulatory mechanisms could contribute to changes in β-glucan and chitin content.

### Chemical inhibition of Gwt1 in several industrially important filamentous fungi has a contrasting effect on macro-morphology in comparison to A. oryzae

We then tested whether the morphology-modulating property of MGX is extendable to other industrially relevant filamentous fungi such as *Aspergillus nidulans*, *Fusarium venenatum* and *Trichoderma reesei.* In contrast to *A. oryzae*, the 3 other fungi exhibited dispersed mycelial growth in their unperturbed states. Intriguingly, addition of MGX converted dispersed hyphae to pellet formation in all the 3 strains, which is completely opposite to the effect of MGX observed in *A. oryzae* (**Figure 6a**). The formation of small compact mycelial pellets reduced the culture viscosity generally (**Figure 6b**), except in 5-day-old MGX-treated *T. reesei* despite the small pellet morphology. These findings demonstrate the potential of MGX application to modulate fungal morphology and reduce culture viscosity across diverse filamentous fungal systems.

**Figure 6:**
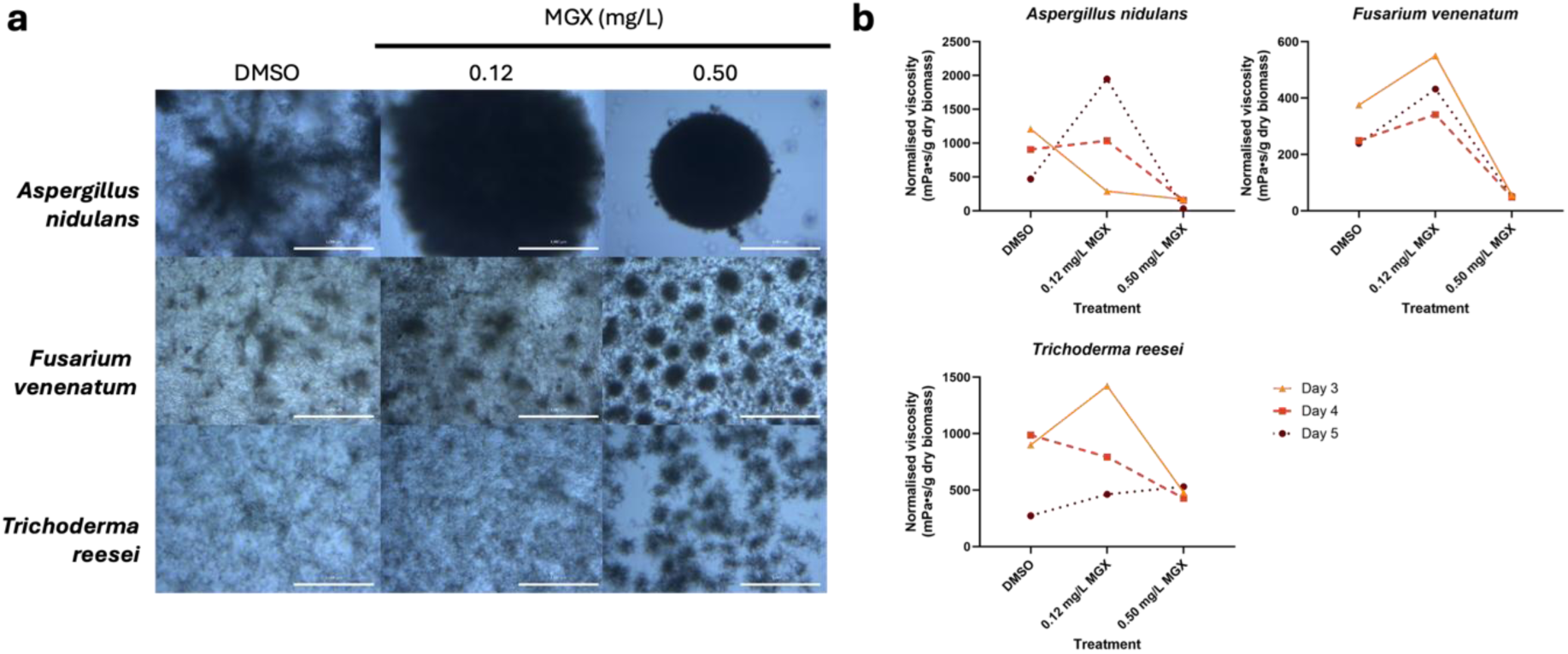
Treatment of other industrially important filamentous fungi favours mycelial pellet formation. (a) Macro-morphology of *A. nidulans*, *F. venenatum* and *T. reesei* cultured in 5×DPY for 5 d. Scale bar, 1000 µm. (b) Biomass-normalised viscosity of *A. nidulans*, *F. venenatum* and *T. reesei* cultured in 5×DPY for 3-5 d.

## Discussion

We provide multiple lines of evidence demonstrating that the disruption of GPI-AP biosynthesis pathway alters the morphology of *A. oryzae* during submerged fermentation. Both the *Δmcd4* and MGX-treated AO1 exhibited hyper-branching and swollen hyphae phenotypes. Chemical perturbation of GPI-anchor biosynthesis pathway via MGX treatment effectively altered macro-morphology from spherical pellets to dispersed mycelia. Alanine-substitution of Gwt1 residues involved in MGX interaction abolished the morphology-modulating effect, confirming that inhibition of Gwt1 function underlies the morphological changes. The ssNMR analysis revealed extensive cell wall remodeling, including redistribution of glucan and chitin components and depletion of mannan and GalN. Transcriptomic profiling reveals distinct adaptive responses to GPI-anchor perturbation-induced cell wall stress in *Δmcd4* and MGX-treated AO1. Repression of cell-cell fusion under GPI-anchor perturbation suggests a potential role in regulating hyphal aggregation and pellet formation. Notably, MGX-induced morphological changes were species-specific, promoting dispersed growth in *A. oryzae* and favouring compact pellet in other filamentous fungi. The transition from pellet to dispersed mycelia enhances biomass accumulation, improving nutrients and oxygen absorption; however, this morphological advantage does not translate into increased protein production. Despite higher biomass in MGX-treated AO1, the biomass-normalised protein secretion was reduced by 30% on day 3, indicating a bottleneck in secretion machinery caused by GPI-anchor biosynthesis disruption. We attribute this to pleiotropic disruption of GPI-anchor-dependent processes, including cell wall assembly and protein trafficking. These findings establish that morphology alone is insufficient to drive protein secretion and underscore the need to decouple morphological control from secretion function. It is therefore critical to identify GPI-anchor function-dependent factors that specifically regulate macro-morphology, to enable precise morphology engineering while preserving protein secretion capacity.

The ssNMR analysis of cell wall composition revealed redistribution of chitin and α-1,3-glucan allomorphs in the rigid phase and a reduction in mannan and GAG in the mobile phase under GPI-anchor-perturbed conditions, with complete depletion of GalN in MGX-treated AO1. While the mechanism underlying polysaccharide allomorph formation remains to be elucidated, GPI-anchor perturbation favoured type-b chitin and type-b α-1,3-glucan allomorphs, likely as a compensatory response to cell wall instability caused by the disruption. Further studies could uncover the mechanisms governing polysaccharide allomorph formation and how these structural changes influence fungal morphology. Depletion of GM is expected, as GPI anchors mediate its transport and covalent attachment to β-glucans in the cell wall ^62^. While mannan levels were reduced, Gal*f* content remained largely unchanged, consistent with its role as a side-chain moiety that does not directly depend on GPI-anchor linkage ^14^. Given that GM is a key structural component that crosslinks with chitin-β-1,3-glucan complex, its depletion is expected to compromise cell wall integrity ^44–47^. We propose that disruption of this chitin-β-1,3-glucan-GM scaffold weakens that cell wall architecture, thereby driving the hyperbranched and swollen hyphal phenotypes observed in GPI-anchor-perturbed strains (**Figure 7**).

**Figure 7:**
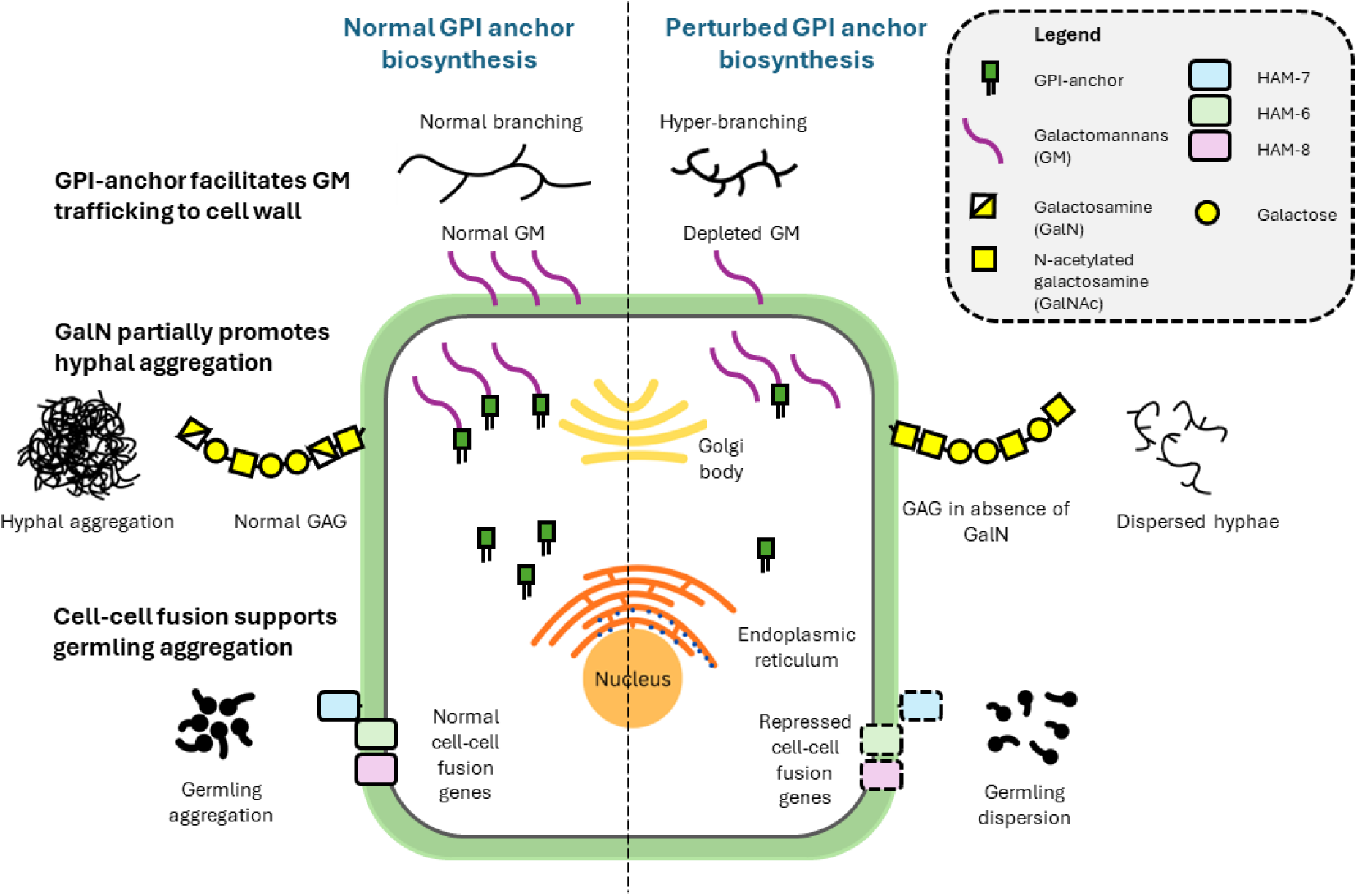
A model to explain how cellular changes in *A. oryzae* AO1 cells subjected to GPI-anchor biosynthesis perturbation cause changes in morphology. GPI-anchor biosynthesis perturbation causes GM depletion, weakens cell wall by reducing GM crosslink with chitin-β-1,3-glucan complex and results in hyper-branched and swollen filaments phenotypes. Chemical perturbation of GPI-anchor biosynthesis via MGX-treatment results in complete depletion of GalN indicating the importance of GalN in promoting hyphal aggregation. GPI-anchor biosynthesis perturbation represses cell-cell fusion genes, including the downregulation of *N. crassa* HAM-7 homolog that encodes a predicted GPI-anchor protein. Cell-cell fusion is hypothesised to promote germling aggregation which may affect fungal macro-morphology.

A feature unique to MGX-treated AO1 was the complete depletion of GalN in the GAG fraction of MGX-treated AO1, a modification thought to implicate hyphal aggregation through hydrogen bonding ^10^. While the reason underlying GalN loss remains unclear, downregulation of *agd3* may contribute to GalN depletion. We propose that the loss of GalN underlies the altered pellet morphology observed in MGX-treated *A. oryzae* (**Figure 7**). Transcriptomics analysis further identified cell-cell fusion pathways under GPI-anchor-perturbed conditions as a key determinant of morphology. Time-resolved MGX addition demonstrated maximal morphological impact at conidial stage (**Figure 2e**), consistent with previous reports on high cell-cell fusion frequency amongst germlings in *N. crassa* ^59^. Notably, a GPI-AP homologous to *N. crassa* HAM-7, which functions with transmembrane proteins HAM-6 and HAM-8 to activate MAK-1 (mitogen-activated protein kinase) pathway during cell-cell signaling ^63,64^, was downregulated. We therefore propose that GPI-anchor perturbation impairs HAM-7 function, disrupting its interaction with HAM-6 and HAM-8, attenuating cell-cell fusion event, thereby preventing hyphal aggregation and pellet formation (**Figure 7**).

Overall, this study highlights the distinct responses of *A. oryzae* to GPI-anchor perturbation under genetic and chemical perturbation conditions, which likely underlie the observed morphological differences during submerged fermentation. The acute cell wall stress induced by MGX treatment triggered extensive transcriptional reprogramming, leading to pleiotropic effects that drove a pronounced morphological transition from pellet growth to dispersed mycelia. To further confirm that partial inhibition of Gwt1 is responsible for the observed morphological change, future studies employing tunable promoter systems to precisely regulate *gwt1* transcription may provide additional mechanistic validation.

Contrary to expectation, MGX treatment of *A. nidulans*, *F. venenatum* and *T. reesei* promoted the formation of small, compact pellets rather than freely dispersed mycelia. While unperturbed *A. oryzae* strains form pellets, these other fungi predominantly form dispersed mycelial network under the same conditions, indicating fundamental differences in cell wall architecture. Such differences likely shape the morphological response to GPI-anchor perturbation. We propose that different fungal species may deploy tailored GPI-anchor protein regulatory programmes to meet their specific cell wall requirements under stress, thereby determining their morphological outcomes. Comparative analyses of cell wall polysaccharide remodelling across fungal systems under GPI-anchor perturbation will be essential to elucidate the mechanistic role of cell wall composition in governing macro-morphology.

## Conclusion

In conclusion, treatment of *A. oryzae* AO1 with MGX effectively converts the compact pellets to freely dispersed mycelia, revealing a tunable morphological state that is highly relevant for industrial fermentation. Our findings underscore the critical role of GPI-anchor-dependent GM trafficking in maintaining cell wall organization, as evidenced by the selective depletion of mannans chains under GPI-anchor-perturbed conditions. Although the mechanism underlying *agd3* downregulation and complete loss of GalN in MGX-treated AO1 remains unknown, this finding suggests that GalN may be responsible for hyphal aggregation. Remarkably, MGX elicits opposite morphological outcome in other industrially relevant fungi, promoting the transition from freely dispersed mycelia to pellet formation. These results highlight the species-specific consequences of GPI-anchor perturbation, while establishing GPI-dependent cellular processes as a viable target for rational engineering of fungal morphology.

## Materials & Methods

### Gene knockout via CRISPR-Cas9

Strains and plasmids used in this study are listed in **Supplementary Table 4**, and primers and sgRNAs are listed in **Supplementary Table 5**. CRISPR/Cas9 gene-knock out was conducted using pHT001 modified from pPTR II plasmid (Takara Bio, Japan), containing yeast auxotrophic marker (TRP1), yeast origin of replication (ARS1), two sets of sgRNA cassette and Cas9 enzyme ^27^. Two crRNAs were selected for each gene of interest using https://crispr.dbcls.jp/ and http://crispor.tefor.net/. Construction and amplification of sgRNA cassettes were conducted via PCR using Q5 High Fidelity DNA polymerase (New England Biolabs, USA) and single-stranded DNA oligonucleotides from Integrated DNA Technologies (IDT, USA). The pHT001 plasmid was digested with SmaI and NotI and the two sets of sgRNA cassettes were inserted into the cut vector through yeast gap repair using *S. cerevisiae* strain (W303 strain). Plasmids were recovered from yeast and amplified by transforming Endura Electrocompetent *Escherichia coli* cells (LSC Biosearch Technologies, United Kingdom). The CRISPR plasmids were purified using Fastfilter Plasmid DNA Maxi Kit (Omega Bio-tek, USA) and transformed into *A. oryzae* via protoplast-mediated transformation.

### Protoplast-mediated Aspergillus transformation

The lysozyme-producing *A. oryzae* strain RIB40 *ΔwA::amyB-*Lys-Arg*-HLY* (AO1) was used as a transformation host. Dextrose-polypeptone medium (DP; 2% dextrin, 1% polypeptone, 0.5% KH_2_PO_4_ and 0.05% MgSO_4_.7H_2_O [pH 5.5]) and potato dextrose agar (PDA; Sigma Aldrich, USA) were used for normal growth and maintenance of all strains. *A. oryzae* was transformed as described previously^65^. Briefly, mycelia cultured overnight in DP media are collected with sterile miracloth (Calbiochem 475855, Merck Millipore, US) and washed with sterile milli Q water. The mycelia were incubated in solution containing Yatalase (Takara Bio, Japan) at 30°C for 3 hours to form protoplasts. CRISPR plasmid (5–10 µg) was added to protoplast suspension (200 µL) that is adjusted to 1 × 10^7^/mL and incubated on ice for 30 min. Poly(ethylene glycol)-containing solution was added to protoplast suspension and incubated at room temperature for 20 min. The mixture was added to Top Agar (21.86% sorbitol, 2% dextrin, 0.8% agar, 0.2% KCl, 0.15% NaNO_3_, 0.1% KH_2_PO_4_, 0.05% MgSO_4_.7H_2_O, 0.002% FeSO_4_.7H_2_O) with 0.1µg/mL pyrithiamine hydrobromide (P0256, Sigma Aldrich, USA) and plated onto Bottom Agar (1.5% agar). Transformation plates were incubated at 30°C for 4 days. The colonies were sub-cultured onto CDex agar (21.86% sorbitol, 2% dextrin, 1.8% agar, 0.3% NaNO_3_, 0.2% KCl, 0.1% KH_2_PO_4_, 0.05% MgSO_4_.7H_2_O, 0.002% FeSO_4_.7H_2_O) with 0.1µg/mL pyrithiamine hydrobromide.

### Genomic DNA extraction for confirmation of gene knock-out

Spores isolated from transformation plate were inoculated into potato dextrose broth (Sigma Aldrich, USA) in screw cap tubes with 1 mm glass beads and incubated at 30°C for 48 hours. The media was then removed, 400 µL of DNA extract solution (0.1M Tris-HCL pH 8.0, 2% Triton X-100, 1% SDS, 10mM EDTA, 0.1M NaCl) and 400 µL of phenol:chloroform:isoamyl alcohol (25:24:1 v/v) was added and the mycelia was lysed in a bead homogeniser at 1,500 rpm for 1 min. Equal volume of isopropanol was added to the top layer in a clean tube to precipitate the DNA, and the pellet was washed with 1 mL of 70% ethanol. The DNA pellet was resuspended in sterile miliQ water and kept at –20°C until use. 2×Taq Mix Red (PCR Biosystems, United Kingdom) was used for colony PCR to confirm gene knock-out.

### Protein sequence alignment and structural alignment

The AoGwt1 (AO090005001245) and ScGwt1 (YJL091C) protein sequences were downloaded from FungiDB release 68 and aligned using Jalview v2.11.5.1 ^66^. The protein structures of AoGwt1 (PDB ID: Q2UQH4) and ScGwt1–MGX complex (PDB ID: 8XIK) were downloaded from AlphaFold Protein Structure Database ^31^ and aligned using PyMOL v3.1.6.1.

### Enzymatic assays

Approximately 5 × 10^5^ conidia in saline Tween solution (0.02% Tween, 0.85% NaCl) were inoculated into 50 mL of 5×DPY (100 g L^-1^ dextrin, 50 g L^-1^ polypeptone, 25 g L^-1^ yeast extract, 5 g L^-1^ K_2_HPO_4_ and 0.5 g L^-1^ MgSO_4_, [pH 8.0]) in a 250 mL Erlenmeyer flask, and incubated at 30°C with agitation at 140 rpm. The lysozyme activity was measured as described by Morsky and Aine ^67^. Briefly, 20 µL of culture supernatant is added to 80 µL of 150 µg/mL suspension of *Micrococcus lysodeikticus* ATCC 4698 lyophilized cells (Sigma-Aldrich, United States) in 50 mM phosphate buffer (pH 6.2). The decrease in absorbance at 450 nm, that was due to lysis of bacterial cells, was monitored at room temperature. The gradient of absorbance over time for culture supernatant was compared to standards containing 2, 1, 0.5, 0.25 and 0 mg/L HLY to determination of HLY concentration.

The amylase activity was measured by quantifying the reducing sugar released by α-amylase using 3,5-dinitrosalicylic acid (DNSA)^68,69^. Briefly, 40 µL of supernatant was added to 160 µL starch solution (1%) and incubated at 37 °C for 10 min. Equal volume of DNSA reagent was added to the reaction mix and boiled at 100 °C for 15 min. The absorbance of the reaction mix at 540 nm was compared to 0 to 10 mM glucose calibration curve and α-amylase activity is reported in 1 U (µmol of glucose released per min).

### Viscosity and biomass measurement

The 5×DPY fungal broth viscosity was measured using viscometer (Atago, Japan), consisting A1 spindle and 15-mL glass beaker, set at 250 rpm and stabilised for 30 s. The dry biomass was measured by filtering 15 mL fungal culture through miracloth (Sigma-Aldrich, USA) to collect the biomass, followed by washing with sterile miliQ water, drying the biomass and freeze-drying the biomass for at least 2 days.

### Cultivation of fungal strains for morphology characterisation

For hyphal morphology analysis, 1 × 10^3^ spores/ mL was cultivated in *Aspergillus* minimal medium (AMM) ^70^ on cover slips in 6-well plates, incubated for 18 h at 30 °C. The cover slip was washed with deionised water and stained with calcofluor white stain (18909, Sigma-Aldrich, United States) before mounting onto glass slide. For pellet morphology, 1 × 10^4^ spores/ mL was cultivated in 5×DPY, incubated for 3 – 4 days at 30 °C, 140 rpm. The mycelia from the 5×DPY culture was diluted by 20× with saline Tween solution and analysed on 6-well plate.

### Microscopy for morphology characterisation of mutants

Hyphal morphology was analysed using Evos FL Bench Top Fluorescent Microscope (Thermo Fisher Scientific, United States) in bright-field and DAPI mode. Macro-morphology was analysed using Zeiss Primovert Inverted microscope (Carl Zeiss AG, Germany) in bright-field and auto-exposure mode. Image analysis was conducted using ImageJ v2.14.0 ^71^ as previously described ^27^. Briefly, the images were converted to 8-bit grey-scale and a grey-scale threshold calculated using Otsu algorithm to obtain a binary image. The macro-morphological parameters such as projected area, Feret’s dimeter, solidity and aspect ratio were measured and calculated from the binary images. Morphology numbers were calculated from these obtained values ^72^.

### Preparation of cultures and ¹³C and ^15^N labelling

Solid cultures of *A. oryzae* AO1 and *Δmcd4* mutant strains were first prepared using non-labelled conditions on a dextrose-based minimal medium supplemented with 1.5% agar. Isotopic labelling was subsequently performed by inoculating to a minimal medium containing 10.0 g/L of ^13^C-glucose and 6.0 g/L of ^15^N-sodium nitrate for uniform incorporation of isotopes into fungal biomass. The media was supplemented with 1 mL/L of trace-element solution (0.04% Na_2_B_4_O_7_·10H_2_O, 5 mM FeCl_3_) and 0.2 M HCl for preventing oxidation. Additionally, a solution containing 0.05% KCl, 0.08% MgSO_4_·7H_2_O, and 0.11% KH_2_PO_4_ was added. The medium was adjusted to pH 6.5. Parallel untreated and drug-treated mycelial cultures were prepared, and 0.5 mg/L MGX was added directly to the medium for the treatment condition. Mycelia were harvested after 3 days of growth. Cultures were grown in 100 mL liquid medium in 250 mL Erlenmeyer flasks at 30 °C with shaking at 210 rpm. Mycelia were collected by centrifugation at 13700 × g for 20 min and washed thoroughly with pure water. Approximately 30 mg of whole-cell material from each sample was packed into a 3.2 mm magic-angle spinning (MAS) rotor for solid-state NMR characterisation.

### Solid-state nuclear magnetic resonance

Solid-state NMR experiments were conducted on a Bruker Avance Neo 800 MHx spectrometer at the Max. T Roger NMR facility (Michigan State University, USA). ^13^C spectra were acquired using a 3.2 mm HCN probe under 15 kHz MAS with approximately 35 mg of sample packed in the rotor and a temperature of 277 K. Chemical shifts were referenced to adamantane (CH_2_ at 38.48 ppm). Radiofrequency field strengths were 71.4 kHz for ^1^H and 50-62.5 kHz for ^13^C. The rigid components of the cell wall were probed using CP-based ^13^C-^13^C CORD experiments^73,74^ with 1 ms CP and 53 ms mixing with 32 scans, while the mobile polysaccharides were characterized using DP-based refocused J-INADEQUATE experiments^75,76^ (2-s recycle delay and 32 scans).

Relative carbohydrate composition was determined based on the previous study ^35,42^ by integrating well-resolved cross-peaks from CORD for rigid and INADEQUATE for mobile spectra using Topsin 4.1.4. Peak intensities were then normalized to obtain relative abundances, and uncertainties were estimated from the variance of the selected cross-peaks (Supplementary Table 2). The image was generated using Origin 2021.

### RNA extraction

Approximately 5 × 10^5^ conidia were inoculated into 50 mL of AMM ^70^ in a 250 mL Erlenmeyer flask, and incubated at 30°C with agitation at 140 rpm for 72 hr. Cell biomass was collected by filtering through miracloth, thoroughly washed with sterile water, flash frozen and ground under liquid nitrogen. RNA was extracted using RNeasy Plant Kit (Qiagen, Germany). Briefly, 100 mg ground biomass was weighed into 2-mL screw cap tubes containing 0.5 mm glass beads and 450 µL of Buffer RLT (Qiagen, USA) supplemented with 0.1% β-mercaptoethanol was added. The biomass was sheared in a homogeniser at 6,000 rpm for 30s, for 5 times with 5 min of incubation on ice in between each shearing. The lysate was transferred to QIAshredder spin column (Qiagen, USA) and RNA was extracted following the manufacturer’s protocol. RNA sequencing was conducted by Novogene (Beijing, China) using NovaSeq X Plus systems (Illumina, USA). Reverse transcription-quantitative PCR (RT-qPCR) was conducted by synthesizing cDNA using QuantiTect Reverse Transcription Kit (Qiagen, Germany) and running the RT-qPCR using KAPA SYBR Fast Master Mix qPCR Kit (Roche, Switzerland) on QuantStudio3 qPCR system (Thermo Fisher Scientific, USA).

### Transcriptomic data pre-processing

Raw paired-end Illumina RNA-seq reads (FASTQ) were first assessed using FastQC v0.11.9 to evaluate per-base sequence quality, GC content, duplication levels, and potential adapter contamination ^77^. Adapter removal and quality trimming were performed using Trim Galore v0.6.6 (on paired-end samples) ^78^, and FastQC for adapter detection and post-trimming quality assessment. Trimmed reads were aligned to the *Aspergillus oryzae* reference genome (ASM18445v3) using HISAT2 v2.2.1 ^79^. Alignment files were processed using SAMtools v1.15: SAM files were converted to BAM format, filtered to retain primary alignments, sorted by genomic coordinate, and indexed for downstream quantification and visualization. Gene-level read counts were generated using featureCounts v2.0.3 by assigning aligned reads to annotated gene features ^80,81^. The resulting raw integer count matrix was used as input for differential gene expression analysis in R using DESeq2 v1.38.0 ^82^. Prior to model fitting, low-support genes were filtered by retaining genes with a total count ≥10 reads across all samples. DESeq2 was used to estimate size factors (normalization), model gene-wise dispersion, and test for differential expression using a negative binomial generalized linear model. log_2_FC estimation on AO1 group was used as the reference condition and expression in the *Δmcd4* and MGX-treated AO1 on days 3 and 4 (target group) were compared against AO1 (reference group); therefore, positive log_2_FC values indicate higher expression in target, whereas negative log_2_FC values indicate lower expression relative to reference. log_2_FC shrinkage was applied to improve effect-size stability.

Multiple testing correction was performed using the Benjamini–Hochberg false discovery rate procedure ^83^, and genes with adjusted p < 0.05 were considered significantly differentially expressed. Differential expression results were annotated using gene features parsed from the *A. oryzae* annotation file using genomic range-based annotation tools, and quality assessments/visuali***s***ations such as MA plots, p-value distributions, and variance-stabilized sample distance heatmaps were generated.

### Transcriptomics data analyses

GO term IDs and gene annotations were obtained from *A. oryzae* Gene Ontology data (FungiDB, release 68), with annotation updated using go-basic.obo v1.2 ^84,85^. DEGs were stratified into upregulated (log_2_FC > 1.5) and downregulated (log_2_FC < −1.5) groups and analysed separately. Over-representation analysis (ORA) was performed using *enricher* function in clusterProfiler v4.14.6 ^86^, with DEGs pvalueCutoff and qvalueCutoff set at 0.05 and 0.2, respectively, and using p-value adjustment by Benjamini-Hochberg procedure. The enriched GO terms and pvalues were extracted from *enricher* and filtered by REVIGO ^87^ using medium cutoff of 0.7 for allowed similarity. The filtered GO term with dispensability values less than 0.5 were selected and visualised using ggplot2 v4.0.1 ^88^. The gene number and p-value are represented by bubble size and colour, respectively, on the bubble plot. Gene-concept network analysis was conducted using *cnetplot* function in clusterProfiler v4.14.6 ^86^ to visualise genes shared between different GO terms.

## Acknowledgments

The authors gratefully acknowledge Prof. Jean-Paul Latgé for his guidance and suggestions in fungal cell wall studies, Prof. Jun-ichi Maruyama for his valuable advice on *Aspergillus oryzae* transformation, and Dr. Naazneen Sofeo and Dr. Miselle Tiana Hengardi for their careful review of the manuscript. C.H.T gratefully acknowledges Prof. Sam Li Yau Fong for his academic guidance during her Ph.D. research.

## Contributions

C.H.T. and P.A. developed the study and designed the experiments. C.H.T. performed molecular cloning, microscopy analysis, enzymatic assays, rheology measurements, RNA extraction and transcriptomics data analysis. I.G. and W.T. performed the solid-state NMR experiments and analyses. S.P.Y. conducted transcriptomics data pre-processing. C.H.T. wrote the manuscript with input and critical feedback from P.A., I.G., W.T. and S.P.Y. P.A. is the corresponding author of the paper. All authors reviewed the paper.

