## Supplementary Information for "Perturbing glycosylphosphatidylinositol (GPI)-anchor biosynthesis alters cell wall architecture and modulates fungal morphology"

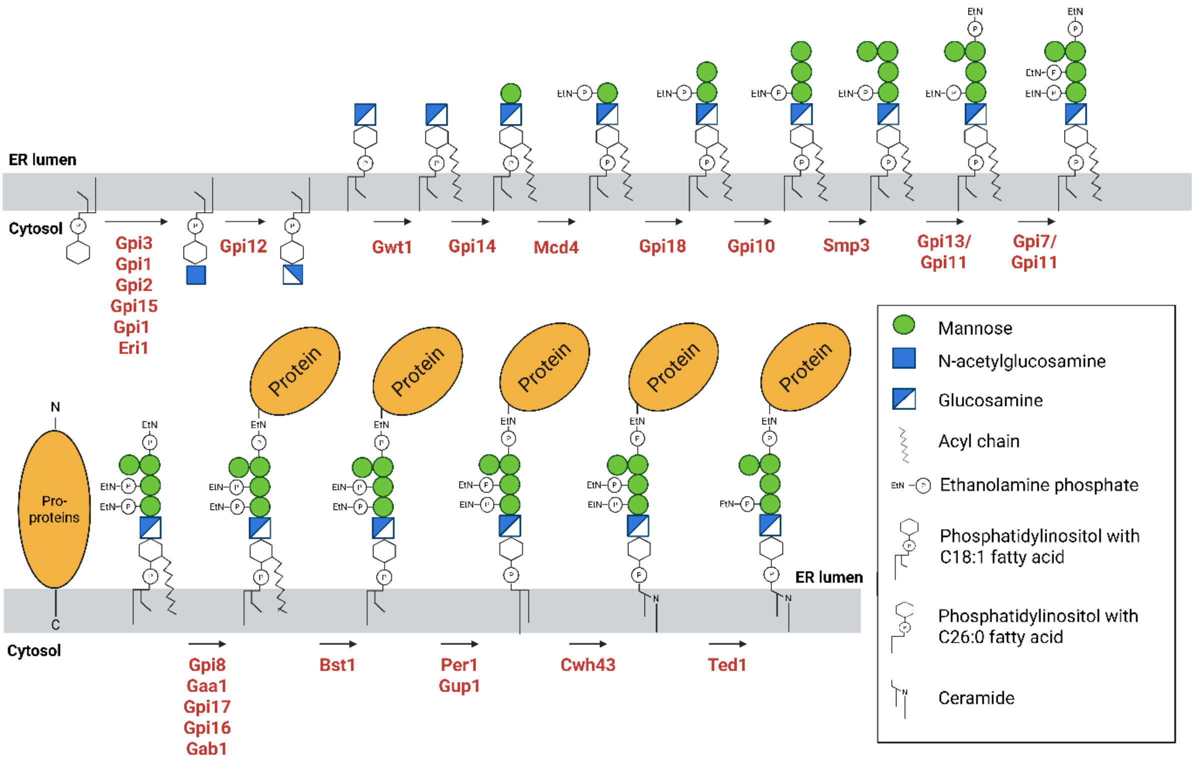


**Supplementary Fig. 1:** Glycosylphosphatidylinositols (GPI)-anchor protein biosynthesis at the endoplasmic reticulum (ER) of yeast and filamentous fungi. Created with BioRender.com.

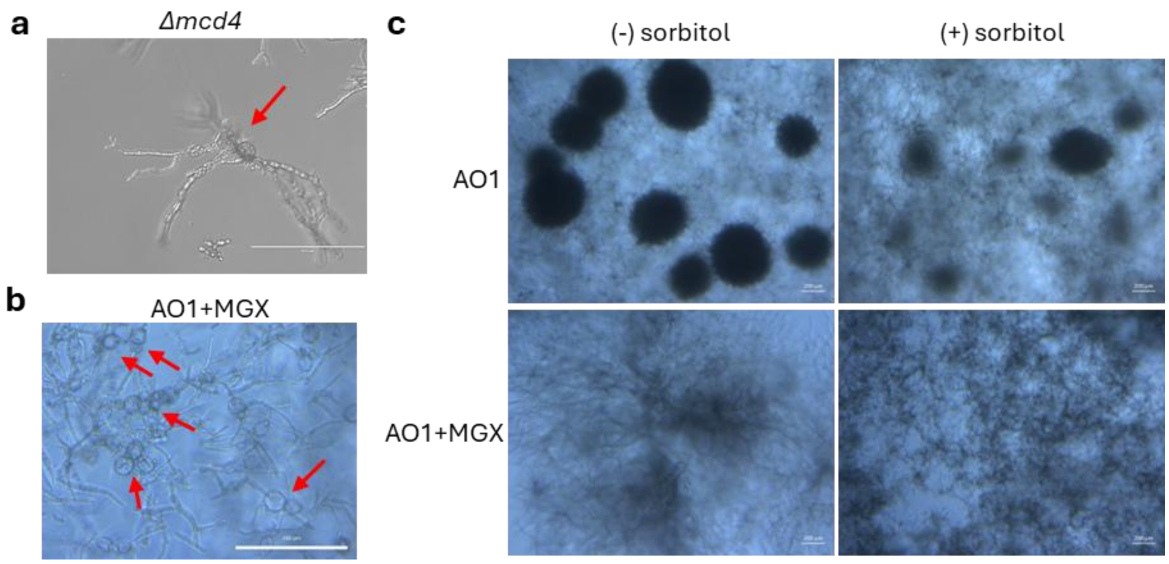


**Supplementary Fig. 2:** Perturbation of GPI-anchor biosynthesis pathway alters morphology of *A. oryzae* AO1. (a) The Δ*mcd4* hyphae exhibit swelling, hyperbranching, and cell wall rupture with release of cytosolic contents (red arrow). Scale bar, 200 µm. (b) MGX treatment induces swollen hyphae in AO1. Scale bar, 1000 µm. (c) Addition of 1.2 M sorbitol to AO1 + 0.5 mg/L MGX did not rescue the pellet morphology after 4 days of growth. Scale bar, 200 µm.

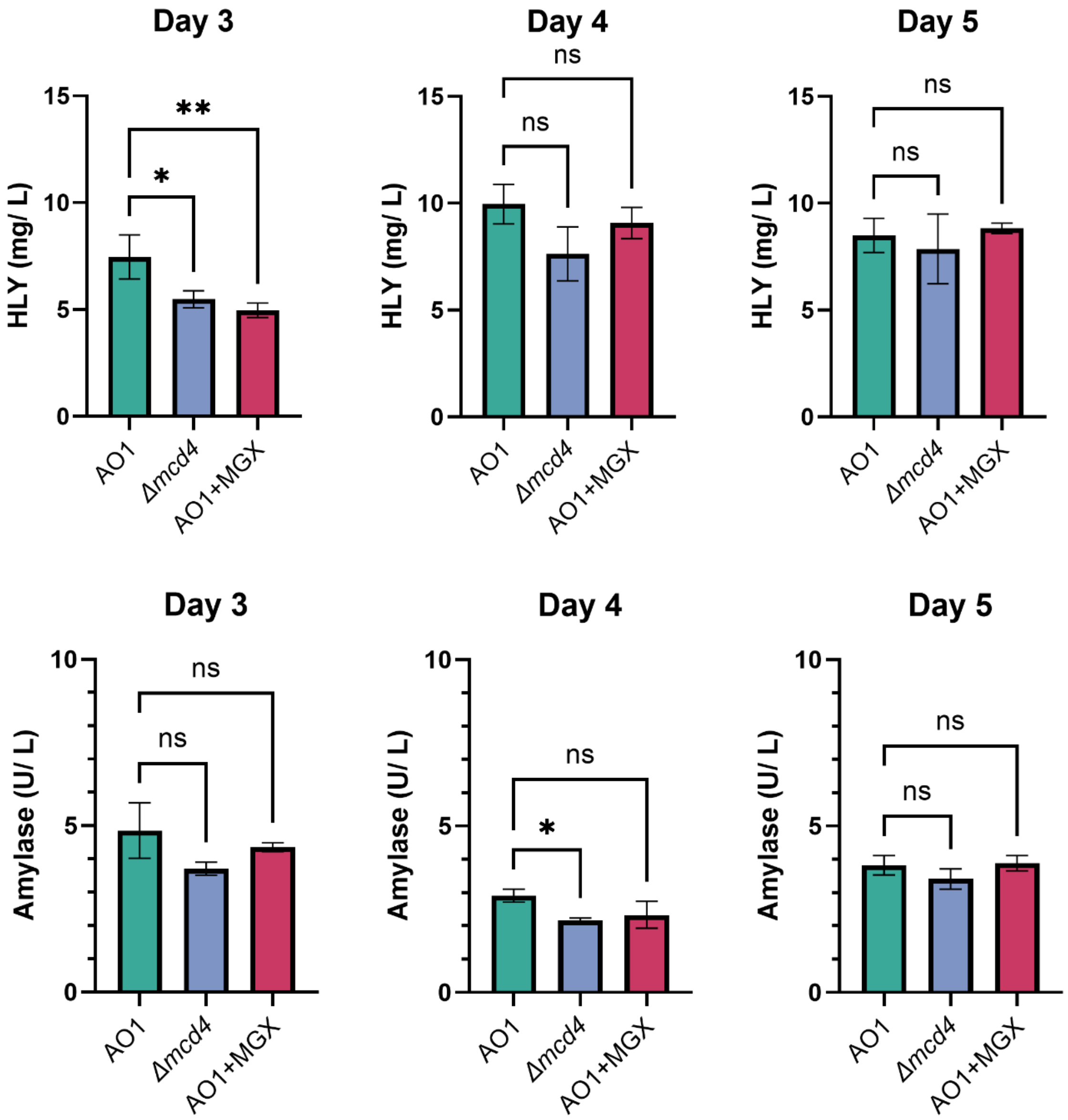


**Supplementary Fig. 3:** Effects of chemical and genetic perturbation of GPI-AP on recombinant (HLY) and native (α-amylase) protein secretion.

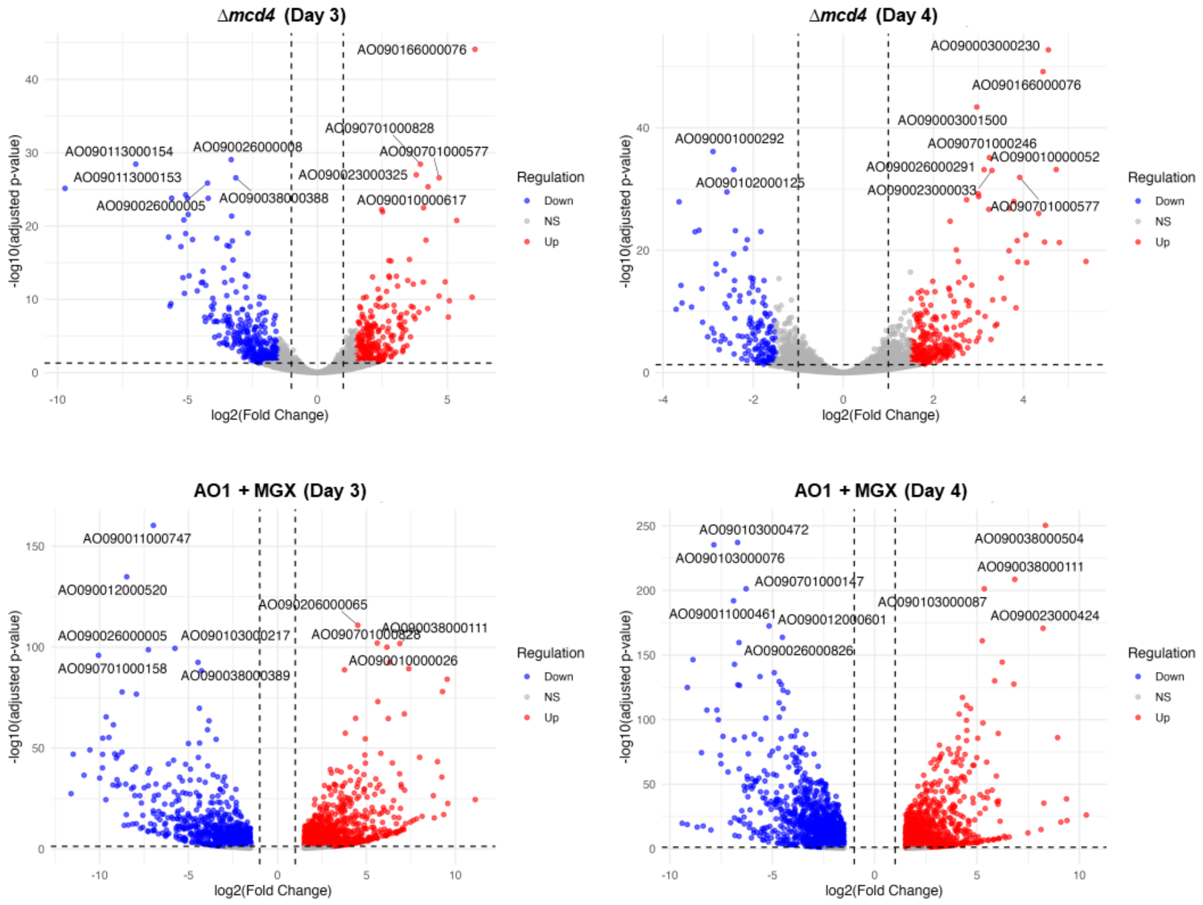


**Supplementary Fig. 4:** Volcano plot of DEGs identified in Δ*mcd4* and MGX-treated AO1 relative to AO1 on days 3 and 4, showing that DEGs in MGX-treated AO1 have a wider range of $\log_2$ fold change values.

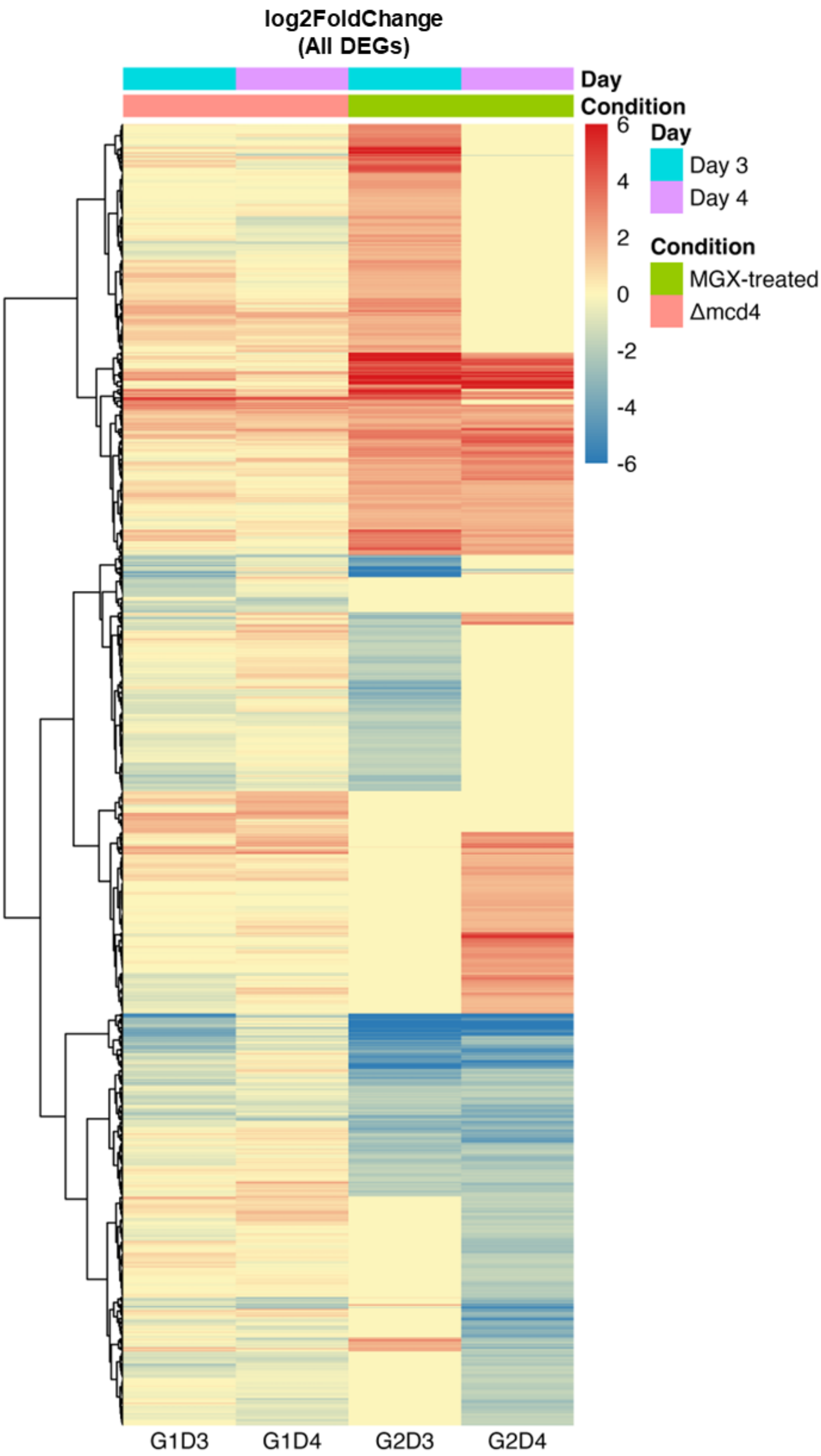


**Supplementary Fig. 5:** Heatmap of DEGs identified in Δ*mcd4* and MGX-treated AO1 relative to AO1 on days 3 and 4, showing distinct DEGs cluster with shared expression patterns across groups and unique expression patterns under specific conditions.

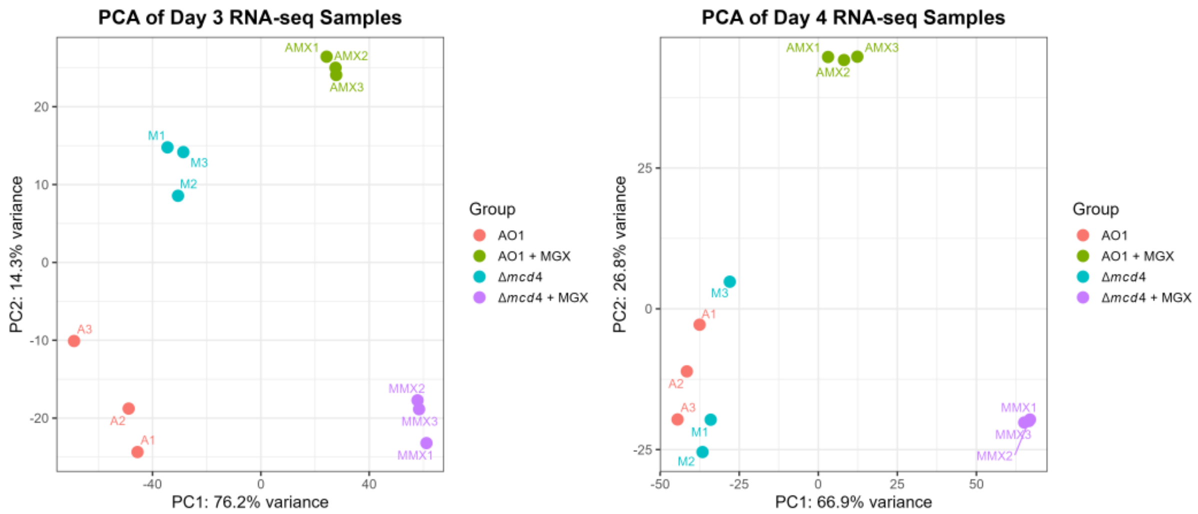


**Supplementary Fig. 6:** Principal component analysis (PCA) showing the clustering of biological triplicates and distinct separation between groups (AO1, AO1 + MGX, Δ*mcd4* and Δ*mcd4* + MGX), except in AO1 and Δ*mcd4* on day 4.

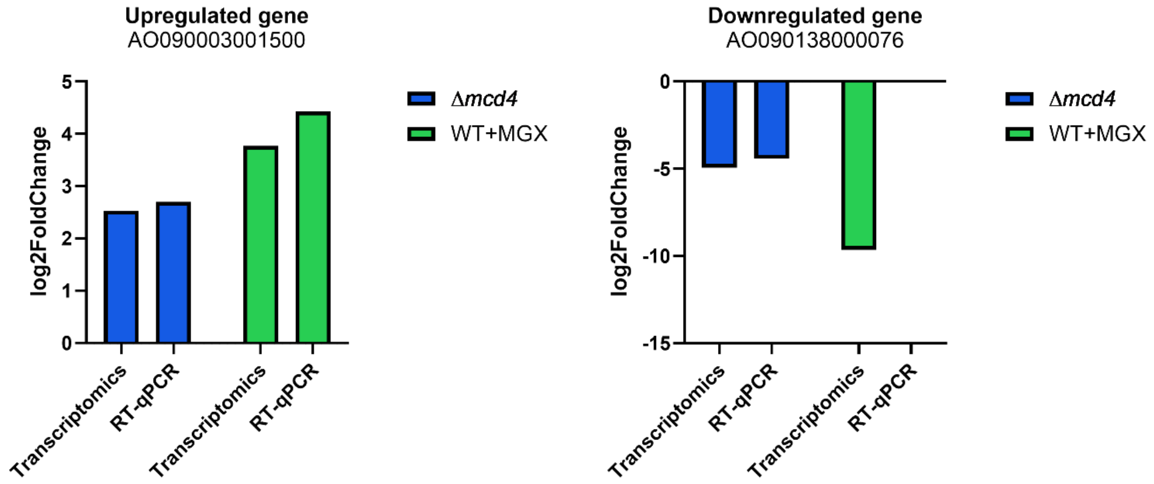


**Supplementary Fig. 7:** Comparison of $\log_2$ fold change values for an upregulated gene encoding α-1,3-glucan/ α-1,4-glucan synthase (AO090003001500) and a downregulated gene encoding stress response protein (AO090138000076) measured by RT-qPCR and RNA-seq (Novogene). RNA samples were collected from Δ*mcd4* and MGX-treated AO1 cultures on day 3.

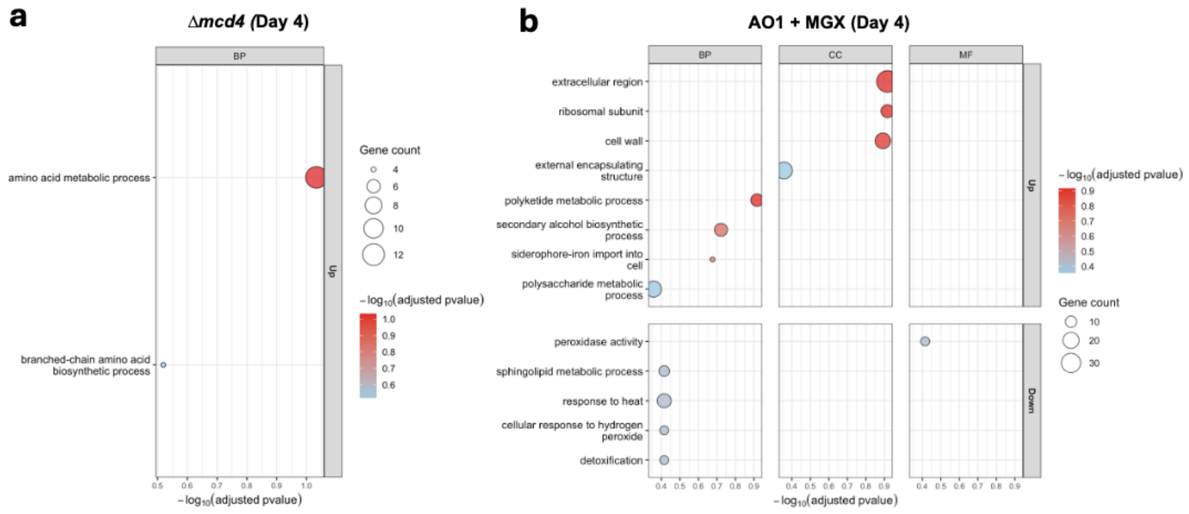


**Supplementary Fig. 8:** GO enrichment plot from over-representation analysis (ORA) conducted on DEGs identified in (a) Δ*mcd4* and (b) MGX-treated AO1 relative to AO1 on day 4. Enriched GO terms were filtered by REVIGO and GO terms with dispensability < 0.5 are displayed in bubble plot. Bubble size represents the gene number while the colour reflects the p-value.

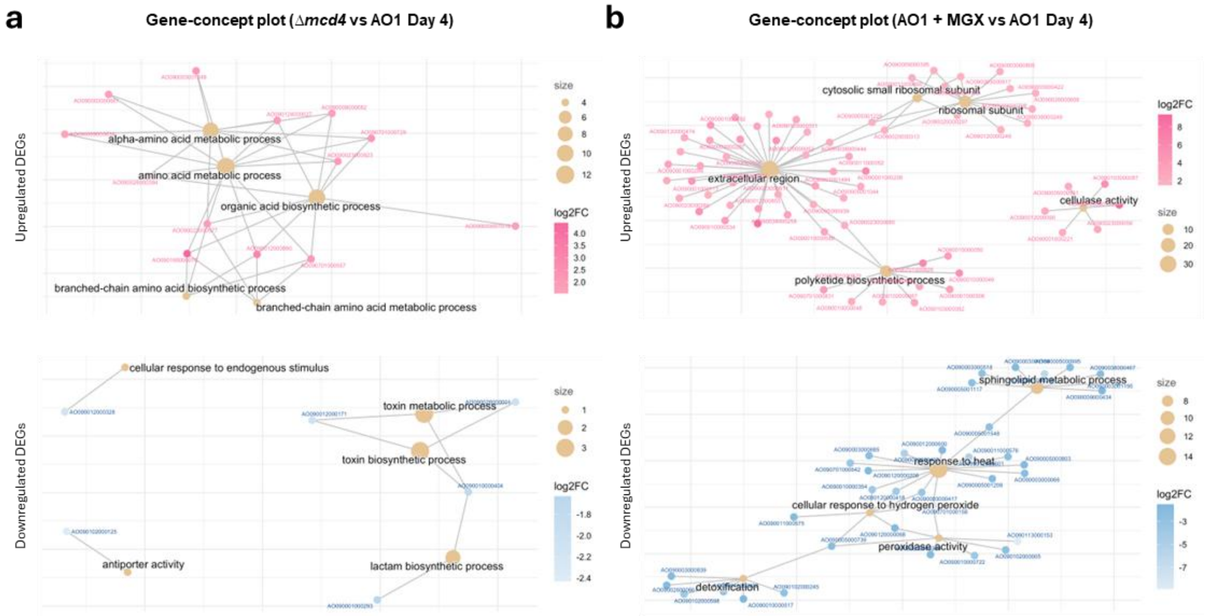


**Supplementary Fig. 9:** Gene–concept networks highlighting DEGs associated with multiple enriched GO terms in (a) *Δmcd4* and (b) MGX-treated AO1 relative to AO1 on day 4. Lines connect shared genes among terms. Each beige node represents a GO term, and node size reflects the number of associated genes. The colour of the gene nodes represents the $log_2FC$.

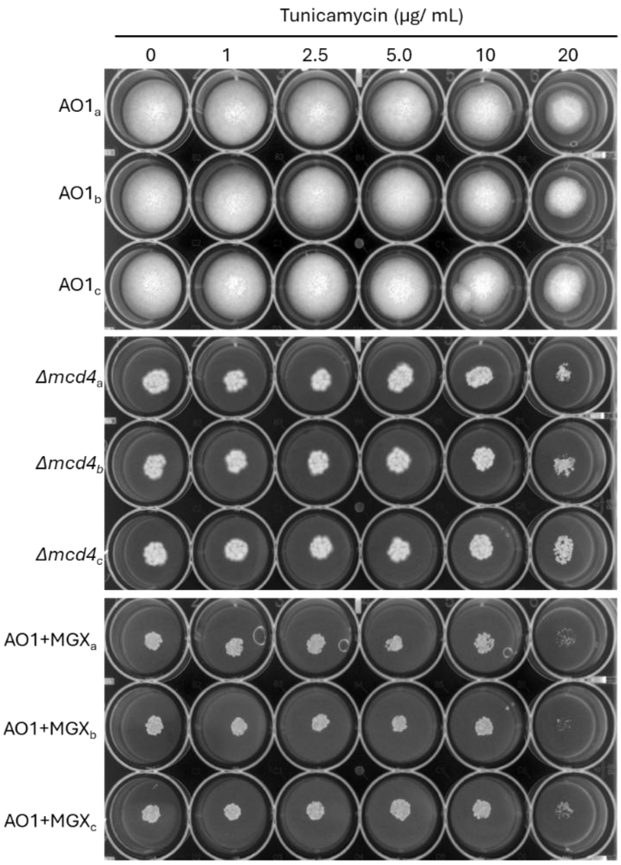


**Supplementary Fig. 10:** Susceptibility of GPI-AP-perturbed strains to ER stress induced by tunicamycin. Conidia ($10^2$) were inoculated in the centre of PDA agar treated with 0 − 20 µg/mL of tunicamycin and incubated at 30°C for 2 days.

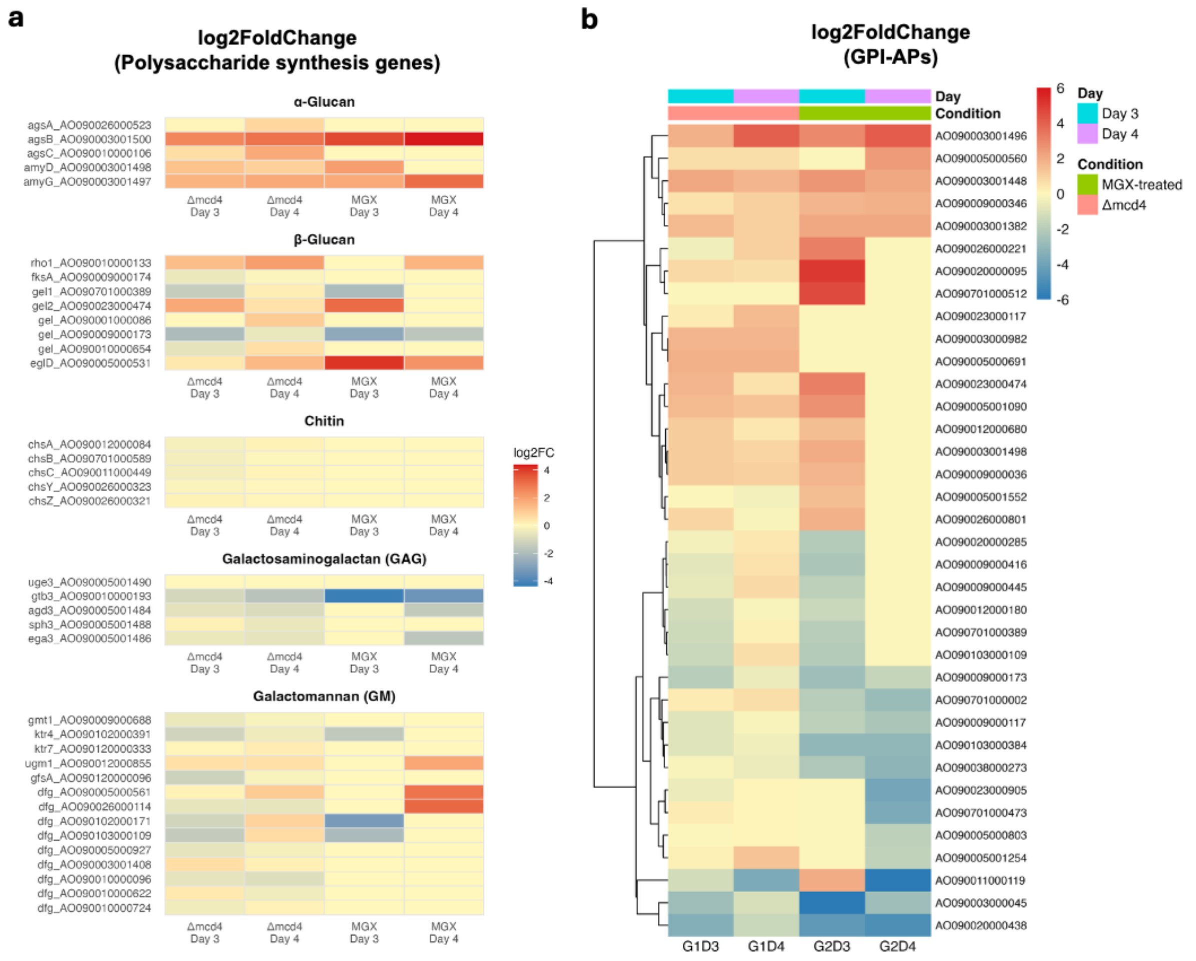


**Supplementary Fig. 11:** Genetic and chemical perturbation of GPI-anchor biosynthesis results in similar gene regulation in genes responsible for polysaccharides synthesis and genes encoding GPI-anchored proteins. (a) Heatmap of $\log_2$ fold change expression of genes involved in α-glucan, β-glucan, chitin, galactosaminogalactan (GAG) and galactomannan (GM) synthesis. (b) Heatmap of $\log_2$ fold change expression of genes predicted to be GPI-anchored proteins.

**Supplementary Table 1:** Predicted GPI-anchored proteins in *A. oryzae* from *in silico* screening of the complete proteome on UniProt and FungiDB using SignalP-6.0 and NetGPI-1.1.

| Gene ID | UniProt ID | Function |
|---|---|---|
| AO090010000654 | Q2TW91 | 1,3-beta-glucanosyltransferase |
| AO090023000474 | Q2UHE9 | |
| AO090001000086 | Q2UP74 | |
| AO090701000389 | Q2U8L0 | |
| AO090009000173 | Q2UUU5 | |
| AO090009000416 | Q2UU86 | Acid phosphatase PHOa |
| AO090003001498 | Q2UIS5 | Alpha-amylase |
| AO090012000663 | Q2UCB4 | Cell wall galactomannoprotein |
| AO090005001552 | Q2UPQ5 | Cellulose-binding GDSL lipase/acylhydrolase |
| AO090701001161 | A0A1S9DR22 | CFEM domain-containing protein |
| AO090023000216 | Q2UI27 | |
| AO090003000045 | Q2UMD2 | |
| AO090103000384 | Q2TY56 | |
| AO090023000239 | Q2UI04 | Chitin deacetylase |
| AO090102000586 | Q2UA11 | Chitinase |
| AO090020000438 | Q2U490 | Copper acquisition factor BIM1-like domain-containing protein |
| AO090120000174 | Q2U6M6 | |
| AO090026000221 | Q2UFG2 | Cupredoxin |
| AO090005000560 | Q2US58 | GH16 domain-containing protein |
| AO090020000289 | Q2U4L7 | Glutaminase A |
| AO090120000279 | Q2U6E0 | Glycosidase |
| AO090005001090 | Q2UQV1 | |
| AO090001000511 | Q2UN44 | |
| AO090005001344 | Q2UQ93 | |
| AO090005000538 | Q2US76 | Glycoside hydrolase family 71 |
| AO090023000915 | Q2UGA5 | GPI anchored cell wall protein |
| AO090010000666 | Q2TW83 | GPI anchored protein |
| AO090011000916 | Q2TZC0 | |
| AO090011000201 | Q2U125 | |
| AO090038000286 | Q2U2X0 | |
| AO090701000382 | Q2U8L4 | |
| AO090012000878 | Q2UBT1 | |
| AO090003001448 | Q2UIW6 | |
| AO090003001326 | Q2UJ78 | |
| AO090003001282 | Q2UJB9 | |
| AO090003000674 | Q2UKU3 | |
| AO090005000610 | Q2US13 | |
| AO090009000445 | Q2UU60 | |
| AO090009000346 | Q2UUE5 | GPI anchored serine-rich protein |
| AO090020000095 | Q2U533 | GPI anchored serine-threonine rich protein |
| AO090012000922 | Q2UBP2 | |
| AO090005000803 | Q2URJ6 | |
| AO090023000905 | Q2UGB5 | GPI-anchored cell wall organization protein Ecm33 |
| AO090003000982 | Q2UK24 | GPI-anchored domain-containing protein |
| AO090011000119 | Q2U193 | Hydrophobic surface binding protein A-domain-containing protein |
| AO090020000588 | Q2U3W7 | Hydrophobin |
| AO090701000473 | Q2U8D2 | Lysophospholipase |
| AO090012000680 | Q2UCA1 | |
| AO090023000685 | Q2UGW4 | |
| AO090003001408 | Q2UJ03 | Mannan endo-1,6-alpha-mannosidase |

| | | |
|---|---|---|
| AO090010000096 | Q2TXL6 | |
| AO090103000109 | Q2TYU3 | |
| AO090009000148<br>AO090005001254 | Q2UUW3<br>Q2UQG8 | Peptidase A1 domain-containing protein |
| AO090012000910 | Q2UBQ4 | PLC-like phosphodiesterase |
| AO090701000002 | Q8NKB6 | Probable aspartic-type endopeptidase opsB |
| AO090023000083 | Q2UIE6 | Probable endo-1,3(4)-beta-glucanase |
| AO090009000117 | Q2UUZ1 | Probable glucan endo-1,3-beta-glucosidase |
| AO090010000729 | Q2TW27 | TSPc domain-containing protein |
| AO090038000622<br>AO090038000273<br>AO090020000279<br>AO090701000512<br>AO090102000201<br>AO090012000664<br>AO090012000359<br>AO090012000180<br>AO090012000013<br>AO090026000801<br>AO090023000117<br>AO090003001496<br>AO090003001382<br>AO090005001359<br>AO090005000691<br>AO090005000652<br>AO090009000467<br>AO090009000036 | Q2U232<br>Q2U2Y1<br>Q2U4M6<br>Q2U8A1<br>Q2UAW9<br>Q2UCB3<br>Q2UD16<br>Q2UDG6<br>Q2UDV5<br>Q2UE13<br>Q2UIC0<br>Q2UIS7<br>Q2UJ25<br>Q2UQ79<br>Q2URU0<br>Q2URX6<br>Q2UU44<br>Q2UV57 | Uncharacterized protein |
| AO090020000285 | Q2U4M0 | WSC domain-containing protein |

**Supplementary Table 2:** Relative molar composition of the rigid and mobile cell wall polysaccharides in *A. oryzae* AO1 and MGX-treated AO1, Δ*mcd4* and MGX-treated Δ*mcd4*. Molar compositions were determined from the integrated intensities of well-resolved signals corresponding to specific carbon sites in 2D $^{13}C$-$^{13}C$ CORD (rigid) and J-mediated INADEQUATE (mobile) components. Values are the average percentages, with errors presenting standard errors derived from spectra analysis. / Represents undetected.

| **Rigid** | **AO1** | **AO1 + MGX** | **Δ*mcd4*** | **Δ*mcd4* + MGX** |
|---|---|---|---|---|
| B | 34±11 | 19±4 | 32±12 | 24±7 |
| G | 2±0.1 | 3±0.1 | 2±0 | 4±0.2 |
| $Ch^a$ | 25±6 | 24±7 | 16±4 | 24±7 |
| $Ch^b$ | 5±0.2 | 16±7 | 13±3 | 16±4 |
| $A^a$ | 21±4 | 21±5 | 27±12 | 18±5 |
| $A^b$ | 8±2 | 14±3 | 7±1 | 11±3 |
| Mn | 5±0.01 | 3±0.1 | 3±0 | 3±0.1 |

| **Mobile** | **AO1** | **AO1 + MGX** | **Δ*mcd4*** | **Δ*mcd4* + MGX** |
|---|---|---|---|---|

| | | | | |
|---|---|---|---|---|
| Gal*f* | 25±11 | 25±8 | 25±9 | 15±3 |
| $Mn^{12}$ | 11±6 | 7±1 | 8±1 | 8±1 |
| $Mn^{16}$ | 13±2 | 3±0 | 9±1 | 9±1 |
| B | 4±0.4 | 14±2 | 20±0 | 9±1 |
| $B^{Br}$ | 15±5 | 15±4 | 18±6 | 15±3 |
| A | 9±1 | 17±5 | 7±1 | 19±5 |
| Gal | 11±1 | 14±3 | 5±1 | 11±2 |
| GalN | 4±0.2 | / | 2±0.1 | / |
| GalNAc | 8±0 | 5±0 | 6±0.5 | 7±1 |
| Ch | / | / | // | 7±0.5 |

**Supplementary Table 3:** Transcription levels of cell-cell fusion genes under GPI-AP-perturbed conditions.

| **Gene name in *Neurospora crassa*** | **Ortholog in *A. oryzae*** | **log2FC** | | | |
|---|---|---|---|---|---|
| | | **Δ*mcd4* vs AO1 Day 3** | **Δ*mcd4* vs AO1 Day 4** | **AO1 + MGX vs AO1 Day 3** | **AO1 + MGX vs AO1 Day 4** |
| *adv-1* | AO090003001259 | -3.8724 | -1.2224 | -5.8027 | -4.5661 |
| *ada-3* | AO090003000967 | 1.6067 | 0.7586 | n.d. | n.d. |
| *so (ham-1)* | AO090003000023 | -1.4770 | -0.8753 | -1.7067 | n.d. |
| *ham-5* | AO090113000103 | -1.7663 | -1.3016 | -1.6390 | -3.8198 |
| *ham-9* | AO090012000944 | -1.5312 | -1.3417 | -2.3966 | -3.5881 |
| *nor-1* | AO090011000671 | -1.2401 | -1.2049 | -1.8762 | -1.9453 |
| *nox-1* | AO090003000460 | -0.6019 | -0.1413 | n.d. | -1.7526 |
| *ham-6* | AO090003000459 | -0.6149 | -0.4799 | -1.6129 | -2.8647 |
| *ham-7* | AO090020000438 | -3.4307 | -1.4840 | -4.5398 | -4.9958 |
| *ham-8* | AO090026000826 | -2.9644 | -1.4804 | -4.1338 | -4.5088 |
| *lfd-2* | AO090026000798 | -1.2996 | -0.7626 | n.d. | -1.5358 |

n.d.: not detected

**Supplementary Table 4: Strains and plasmids used in this work**

| **Strains/ Plasmids** | **Description** | **Source** |
|---|---|---|
| ***A. oryzae*** | | |
| AO1 | RIB40 Δ*wA*::amyB-Lys-Arg-HLY | [27] |
| Δ*mcd4* | *mcd4* deletion in AO1 | This study |
| Δ*agd3* | *agd3* deletion in AO1 | This study |
| **Plasmids** | | |
| pHT001 | *E. coli-Aspergillus* shuttle vector (Takara pPTR II) containing Cas9, ARS1 and TRP1 | [27] |
| pHT001-Δ*mcd4* | pHT001 plasmid containing two sgRNA expression cassettes targeting *mcd4* for CRISPR/Cas9-mediated deletion | This study |
| pHT001-Δ*agd3* | pHT001 plasmid containing two sgRNA expression cassettes targeting *agd3* for CRISPR/Cas9-mediated deletion | This study |

**Supplementary Table 5: Primers and sgRNA used in this work**

| Primer | Sequence | Usage |
|---|---|---|
| HT157 | CTAAAAC**ACACGCGAGTTCCCACCGAA**ACTTGTTCT TCTTTACAATGATTTATTTACCC | Reverse primer for *mcd4* sgRNA1 |
| HT158 | AACAAGT**TTCGGTGGGAACTCGCGTGT**GTTTTAGA GCTAGAAATAGCAAGTTAAAATAAG | Forward primer for *mcd4* sgRNA1 |
| HT159 | CTAAAAC**CGTGTGTGTACCTGGGACTG**ACTTGTTCT TCTTTACAATGATTTATTTACCC | Reverse primer for *mcd4* sgRNA2 |
| HT160 | AACAAGT**CAGTCCCAGGTACACACACG**GTTTTAGA GCTAGAAATAGCAAGTTAAAATAAG | Forward primer for *mcd4* sgRNA2 |
| HT254 | GCAGTGTAACGGCGTATCATCCCTGG | Forward primer to check for *mcd4* deletion |
| HT255 | GGGCTTCCATACCTGCGCACCCATA | Reverse primer to check for *mcd4* deletion |
| HT468 | CTAAAAC**GTGTCAGTACGACAGGCCTC**ACTTGTTCT TCTTTACAATGATTTATTTACCC | Reverse primer for *agd3* sgRNA1 |
| HT469 | AACAAGT**GAGGCCTGTCGTACTGACAC**GTTTTAGA GCTAGAAATAGCAAGTTAAAATAAG | Forward primer for *agd3* sgRNA1 |
| HT470 | CTAAAAC**TTGGCCAGTCTCACCGTTGA**ACTTGTTCT TCTTTACAATGATTTATTTACCC | Reverse primer for *agd3* sgRNA2 |
| HT471 | AACAAGT**TCAACGGTGAGACTGGCCAA**GTTTTAGA GCTAGAAATAGCAAGTTAAAATAAG | Forward primer for *agd3* sgRNA2 |
| HT486 | TCCACGATGCTGACTAGGGA | Forward primer to check for *agd3* deletion |
| HT487 | GTGGTCTAGGCCGGTCATTC | Reverse primer to check for *agd3* deletion |

Note: sgRNA sites are in **bold**.
